# Beyond winglets: evolutionary scaling of flight-related morphology in stick insects (Phasmatodea)

**DOI:** 10.1101/2022.06.09.495408

**Authors:** Yu Zeng, Sehoon Park, Camille Gonzales, Stephanie Yom, Faszly Rahim, Robert Dudley

## Abstract

The first winged insects evolved from a wingless ancestor, but details of the transition to a fully-winged morphology remain unclear. Studying extant pterygotes with partial wings, such as the stick insects (Order Phasmatodea), may help us to understand such a transition. Here, we address how a series of flight-related morphological parameters may correlate with flight evolution by studying different phasmids representing a volancy continuum ranging from miniaturized to full-sized wings. Variation in phasmid wing shape, venation, wing mass and the mass of flight muscle can be described by specific scaling laws referenced to wing length and wing loading. Also, the mass distribution of the body-leg system is conserved in spite of a wide range of variation in body shape. With reduced wing size and increased wing loading, the longitudinal position of the wing-bearing thoracic segments is shifted closer to the insects’ center of body mass. These results demonstrate complex reconfiguration of the flight system during wing morphological transitions in phasmids, with various anatomical features potentially correlated with reduced flight performance attained with partial wings. Although these data represent phasmid-specific features of the flight apparatus and body plan, the associated scaling relationships can provide insight into functionality of intermediate conditions between wingless and fully-winged insects more generally.

## 1. Introduction

Deriving from a single origin within the small and wingless terrestrial hexapods, the winged insects (Subclass Pterygota) and their wings continue to exhibit rich dynamics on evolutionary timescales. The earliest fossils of winged insects derive from the Upper Carboniferous (∼330 million years ago), but intermediates between the wingless apterygote ancestors and their fully-winged descendants remain undocumented, thus posing a major challenge to understanding the origin of insect flight (**Fig. 1A,B**) (Alexander, 2018). The ancestors of winged insects were presumably small hexapods possessing only rudimentary protowings, and subsequently enlarged the wings over a relatively rapid period (Kukalová-Peck, 1983; Schachat et al., 2023). Knowing the morphology of these transitional forms is critical to reconstructing the origins of insect flight, but these forms are as yet undescribed from the fossil record.

**Figure 1.**
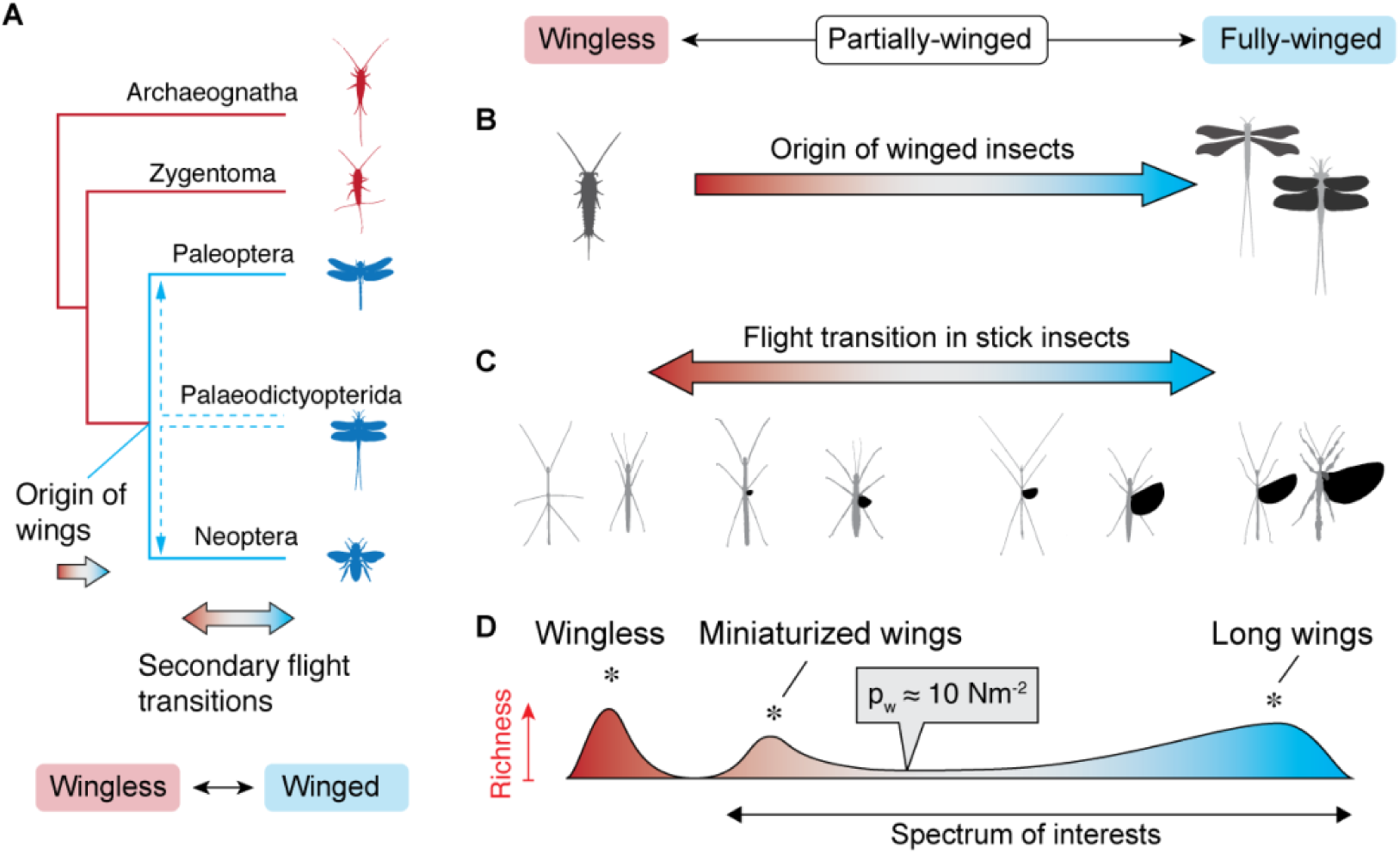
Flight transitions in ancestral pterygotes and phasmids. **(A)** A consensus cladogram of major insect clades, highlighting the initial origin of flight and secondary flight transitions among pterygotes. The phylogenetic position of early pterygote clade Palaeodictyopterida remains unresolved (Prokop and Engel, 2019). **(B)** The initial flight gain that gave rise to the common ancestor of winged insects (Pterygota) was characterized by a transition from wingless to winged forms, but the morphology of partial-winged intermediate stages remains unclear. **(C)** Secondary flight transitions in phasmids are characterized by gains and losses of flight, evident from various sized partial-wings found in different species and sexes. **(D)** Species richness of different phasmid taxa features three peaks with respect to relative wing size and wing loading, which likely correspond to three adaptive peaks (as indicated by asterisks; see Zeng et al., 2020). The current study focuses on the morphological regime lying between long- and miniaturized wings.

How might we quantitatively index morphological intermediates between wingless and fully-winged forms? The most intuitive answer would be to use the size of the wings relative to the body length (henceforth denoted Q), and then the gain of flight may then be correlated with its progressive increase . Nonetheless, powered flapping flight is concurrently underpinned by the generation of sufficient aerodynamic force to offset body weight, at first only in part but progressively increasing in magnitude over evolutionary time. From a biomechanical perspective, studying evolutionary intermediates to volant and wingless forms (here termed ‘flight transition’) must also incorporate variation in wing loading (p_w_, the ratio of body weight to sustaining wing area) and body length (L). For example, the earliest winged insects likely underwent substantial increases in body size (from an initial 1 – 2 cm) while acquiring flapping flight (Ellington, 1991; Wootton and Kukalová-Peck, 2000; Dudley, 2000; **Fig. 1B**). Also, as flight transition may involve interactions between various selective forces acting on both wing and body size (Zeng et al., 2020), it is necessary to discretize flight transition as reconfigurations of flight-related traits within a multidimensional space, with the principal variables minimally including Q, L and p_w_.

Partial-winged insects undergoing flight transition may help to understand the factors influencing the origin of flight. The stick insects (Order Phasmatodea) exhibit vestigial forewings (in most species) and variably sized hindwings over the full range of Q values, from fully-winged (Q ∼0.8) to complete winglessness (Q = 0) (**Fig. 1C**). The size of phasmid wings (and presumably flight capability) show complex phylogenetic patterns, and recent work suggests both flight gains and losses (Bank et al., 2021; Bank and Bradler, 2022; Forni et al., 2022). In general, relaxed selection for flight may underpin various partial-winged morphologies within an arboreal context. Current evidence suggests two ‘adaptive peaks’ associated with either large or miniaturized wings, which are evolutionary consequences of both gains and reductions of flight over various evolutionary timescales (**Fig. 1D**) (Zeng et al., 2020; Forni et al., 2022).

Thus far, flight kinematics has only been studied in a single species of phasmid (Boisseau et al., 2022). For a preliminary assessment of the correlation between flight capacity and wing size, we made highspeed video films of flight in phasmid taxa with different sized wings (see **Methods**). The character of flight trajectories was correlated with wing size: long-winged phasmids were capable of ascending flight, while partial-winged phasmids only performed either gliding or parachuting (**Fig. 2A-C**). Partial-winged phasmids were not able to generate substantial wing forces, and thus relied more on aerodynamic forces generated on the body and legs (which are less efficient aerodynamically) to achieve controlled gliding and parachuting. Also, flight transitions may involve variation in the coordination between wings and the body-leg system, which involves phasic interaction between aerodynamic and inertial forces among these structures. For example, long-winged phasmids flapped wings at high amplitude and oscillated the abdomens in synchrony with wingbeats, whereas short-winged phasmids showed either low-amplitude or no flapping and maintained a relatively stationary abdomen and leg postures during gliding or parachuting (**Fig. 2A-C**). Our first step toward understanding transitions in phasmid flight biomechanics is to address the variation of flight-related morphological parameters with respect to aforementioned principal variables (**Fig. 3**).

**Figure 2.**
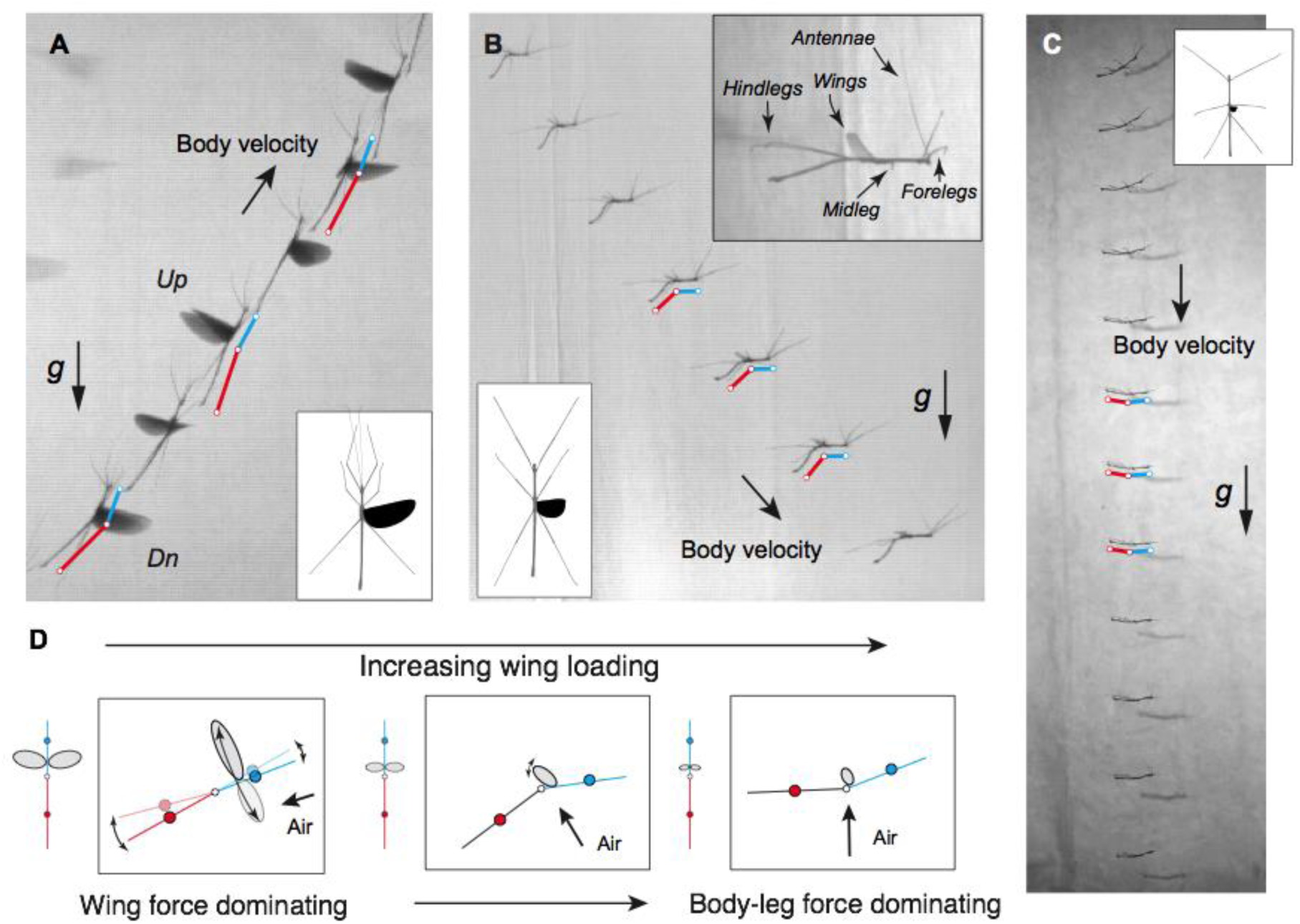
Wing motion and body-leg posture in the flight of different phasmids. Preliminary observations suggest that flight capability is correlated with wing size and wing loading in phasmids. Sample flight sequences for **(A)** *Diardia signata* male (L ∼12 cm; p_w_ ∼3.8 Nm^-2^) in ascending flight, **(B)** *Lopaphus iolas* male (L ∼7.5 cm; p_w_ ∼7.3 Nm^-2^) in gliding flight, and **(C)** *A. tanarata tanarata* female (L ∼3.9 cm; p_w_ ∼30 Nm^-2^) in parachuting. A two-link skeletal model of the body is annotated for each example. Note the oscillation of the abdomen in the *D. signata* male (with ventral flexion during the upstroke [Up] and dorsal flexion during the downstroke [Dn]), which is not evident in partial-winged species. **(D)** Schematics of wing motion and body posture in flights with different sized wings and correspondingly a substantial increase in wing loading, with variable configurations among wings, body, and airflow.

**Figure 3.**
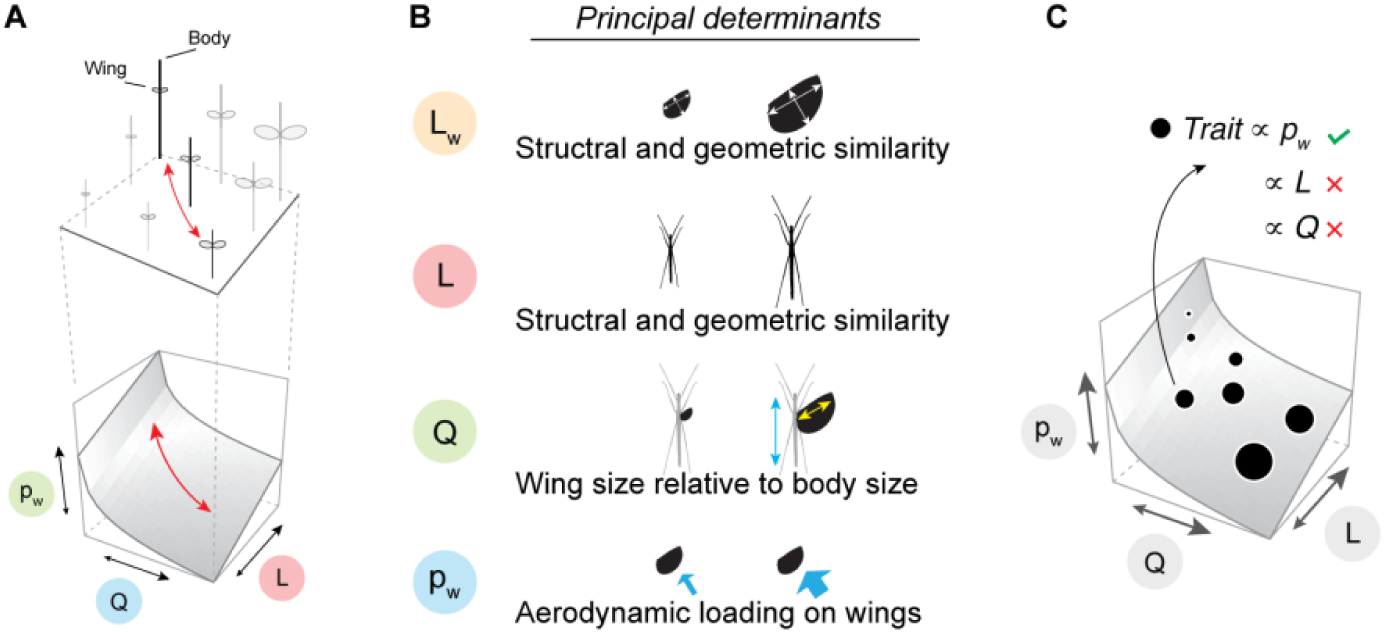
Principal dimensions in phasmid flight transition. **(A)** The aerodynamic capability of a phasmid can broadly be represented by wing loading (p_w_), which scales with relative wing size (Q) and body size (L). Wing loading can then be visualized as a surface (namely the ‘wing loading landscape’), which represents a morphotypes as possible combinations of L and Q. Previous work has quantitatively described the topology of the p_w_ landscape, and identified a general inverse correlation between L and Q (red trajectory; see Zeng et al., 2020), but how other flight-related traits vary along this trajectory remains unclear. See more details in **SI Fig. S6** and **SI Section A**. **(B)** Principal determinants of the four principal variables used in scaling analyses. **(C)** Significant correlations with p_w_ can be used to identify traits influenced by the gradient of flight-related selection in winged phasmids (corresponding to the spectrum of interest in Fig. 1D). For example, the value of a given trait (represented by the size of black dots) may correlate with p_w_ and thus vary along the gradient of p_w_; however, it is not necessarily correlated with L or Q, given that both L and Q may evolve under the influence of numerous non-fight-related selective pressures.

What we know about aerodynamically functional wings mostly pertains to those used in powered flapping flight; these wings are light, tough, and flexible, and repetitively to generate cyclical aerodynamic forces through the wingbeat. Although it is trivial to assume that partial-wings are but reduced versions of full-sized wings, this assumption should be tested because partial-wings may interact with relative airflow in very different ways. Here, we discuss four important features of phasmid wings:

1. Wing shape. The shape of the wing planform is fundamental to assessing the aerodynamics of flight (Ellington, 1984), and variation in wing shape (most broadly represented by the aspect ratio) is specific to insect lineages and relevant to aerodynamic efficiency (Bhat et al., 2019).
2. Wing venation. The venational network, essentially desiccated tracheal tubes, provides structural support for the wing membrane and allows for deformations during flight (Wootton 1992). Compared to wings of close relatives such as the Orthoptera (see Wootton et al., 2000), phasmid wings are uniquely characterized by a narrow, elytrized costal edges and an entire membrane supported by only two radially organized vein groups (**Fig. 4A,B**). The configuration of these veins (e.g., number and hierarchical organization) should inform as to wing deformability in flight, and along with variable developmental pathways relative to wing size.
3. Wing mass. The mass of wings influences inertial power (for accelerating wings) expenditure during repetitive flapping, and thus contributes to flight energetics (Dudley, 2000).
4. Mass of flight musculature. Total flight muscle typically correlates with flight capability and is often evaluated in the form of relative mass, as expressed relative to body mass (Marden, 1989).

**Figure 4.**
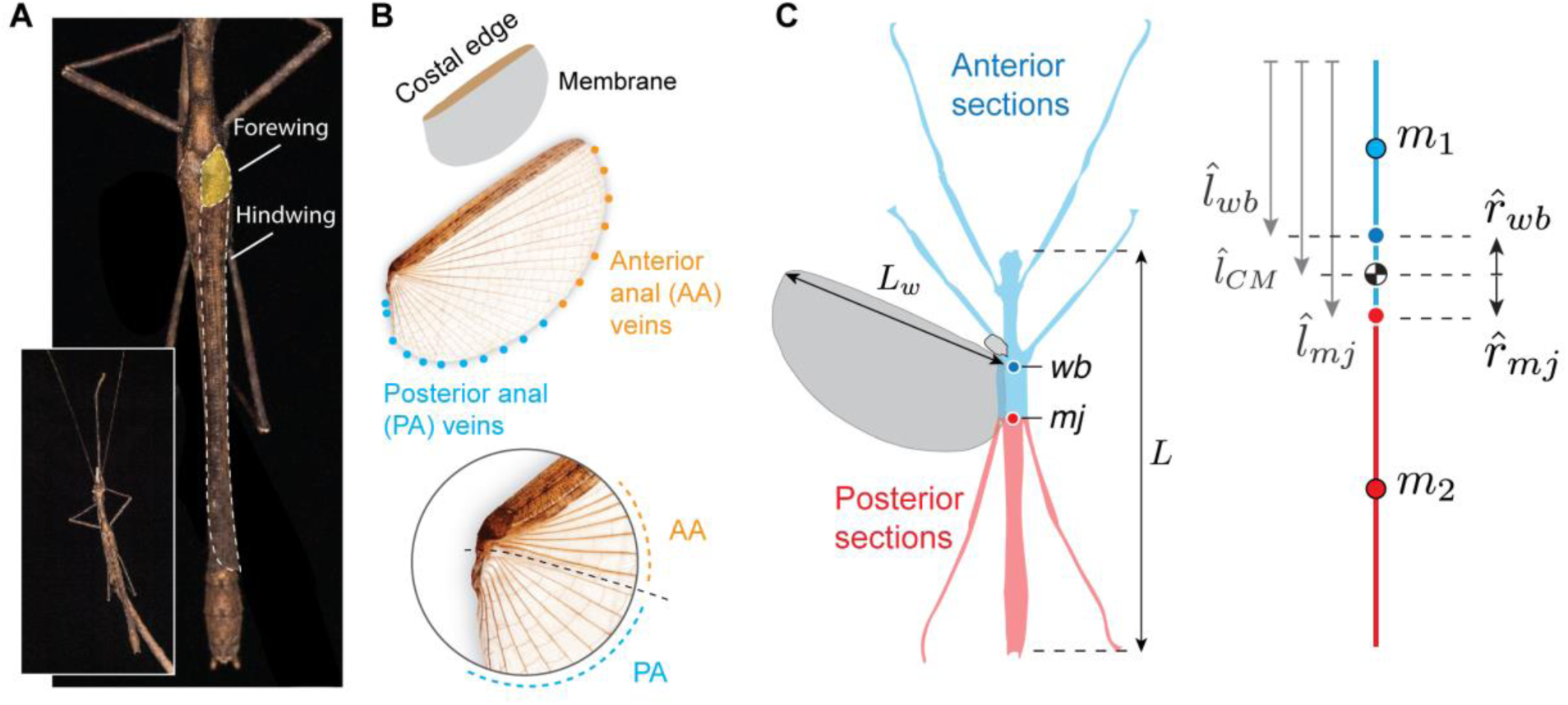
Morphometrics of winged phasmids, demonstrated with a male *Asceles tanarata singapura*. **(A)** When the insect is resting, hindwings are folded with the elytrized costal edges covering the abdomen. The small, elytrized forewings cover the base of hindwings. **(B)** An overview of hindwing components, including the elytrized costal edge and a large membrane supported by two groups of radially arranged anal veins (see **SI Fig. S2** for more details). **(C)** Dorsally projected planform of the insect. Accounting for oscillation about the thorax-abdomen junction at the median joint (mj), the body-leg system can be modeled as a two-link system: an anterior section with mass 𝑚_1_ (blue) that includes the head, thorax and fore- and midlegs, and a posterior section with mass 𝑚_2_ (red) that includes abdomen and hindlegs (see main text). (Right) Configuration for the locations of morphological landmarks, where *l̂_wb_*, *l̂_CM_* and *l̂_mj_* are locations of wingbase (wb), center of mass (CM) and median joint, respectively; *r̂_wb_* and *r̂_mj_* are radii from CM to wingbase and median joint, respectively (see Methods).

We also examine several flight-related parameters of the body-leg system in phasmids, given their slender shapes and flight posture with fully extended legs (**Fig. 4C**).

1. Planform area and shape. The aerodynamic forces on body-leg sections mainly constitute payload drag in flying insects but may play significant roles if the phasmid is gliding or parachuting (Dudley et al, 2007; Zeng et al., 2017). Aerodynamic evaluation requires knowledge of the planform area of body-leg sections; in particular, if simplified as cylinders, the slenderness of these sections can be used to estimate aerodynamic coefficients (Ellington, 1991).
2. Mass distribution. The location of the center of mass CM from head (*l̂_CM_*) is fundamental to biomechanical analysis at the whole-insect scale. For phasmids in particular, mass distribution anterior and posterior to the CM should also be quantified in order to evaluate the inertial dynamics of abdominal oscillation. Such oscillations at wingbeat scale are different from those in hovering fight (e.g., of hawkmoths; Dyhr et al., 2013), and may be more comparable to large abdominal oscillations in butterflies (see Chang et al., 2020).
3. Location of wingbase. The radius from CM to wingbase (*r̂_mj_*) determines how the wing forces may exert rotational moments on the insect’s body (Ellington, 1984).
4. Location of thorax-abdomen junction (in phasmids, the median joint; **Fig. 4C**). When inertial moments are generated by an oscillating abdomen, the radius from CM to the abdomen’s rotational axis (𝑟_𝑚𝑗_) partly determines the moment arm of abdominal mass and correspondingly the magnitude of inertial moment.

Here, we aggregate and analyze morphological data from different phasmid taxa representing a broad range of wing loading (1.4 – 2300 Nm^-2^; **SI Fig. S1**). First, we provide an overview of flight-related morphology in phasmids, as compared to other winged insects, including some of the earliest fossil pterygotes and also eight modern taxa commonly studied for flight biomechanics. Second, we develop linear models to describe various morphological intermediates in phasmid flight transitions. For wing-related variables, we analyzed scaling with respect to L_w_, Q and p_w_. Lacking knowledge on how wing morphology varies through partial-winged morphs that glide or parachute, we instead adopted null hypotheses assuming geometric similarity and structural conservation of partial wings in relation to full-sized wings, such that wing shape and venation are invariant across L_w_ and wing mass would scale with the second or third power of L_w_ (see **Results**). The scaling of body-leg traits was analyzed against L, Q and p_w_; we adopted null hypotheses assuming conserved geometry (i.e., isometry) and significant positive scaling with body length L. For each variable, we derived power-law scaling models using generalized linear regressions, and tested correlations using phylogenetically generalized least-squared (PGLS) models. A significant correlation with wing loading would imply potential selective influence related to the insect’s aerodynamic capability (**Fig. 3A**). In particular, although p_w_ depends on both L and Q, a trait that is linearly correlated with p_w_ may not necessarily be so correlated with either Q or L (**Fig. 3B**). Sensitivity analyses were conducted to address possible stepwise correlations with respect to p_w_, given a possible disjunction of wing function regimes at p_w_ ≈ 10 Nm^-2^ (**Fig. 1D**; Zeng et al., 2020).

Because part of our aim was to identify the influence of flight-related selection, we mainly focused on phasmids with expressed wings (**Fig. 1D**). Our analyses will thus reveal how parameters are correlated along the flight evolution trajectory defined by Q, L and p_w_ regardless of the direction of transition (i.e., gain vs. loss), because the history of wing size and the direction of flight evolution in modern phasmids are largely unclear except for a few taxa. Recognizing this complexity, we prioritized maximizing coverage of wing and body sizes when sampling taxa; nevertheless, we covered one of the few well-documented cases of environment-mediated flight reduction in a single species (details in **SI Section A**).

## 2. Materials and methods

### 2.1 Insect sampling

Our sampling covered field-collected and captive-reared phasmids from five families, but mostly from the tribe Necrosciinae. Our sampling also included the tropical stick insect *Asceles tanarata* Brock, 1999 (Brock, 1999; Seow-Choen, 2000), which consists of three subspecies distributed along an altitudinal gradient, representing one of the few well-documented cases of wing reduction across an environmental gradient (**SI Section A**).

Given sexual dimorphism, male and female insects of the same species were treated as different evolutionary morphs. For convenience, we designate any given sex of a given species to be a ‘sample’ of a specific flight morph. For samples with more than one specimen, the mean value among individuals was used. Morphological data were organized in five main datasets (see workflow, sample sizes, and coverage of wing loading in **SI Fig. S1**). All insects were weighed with a portable balance (PP-2060D, Acculab; accuracy of 0.001 g and linearity of ±0.004 g) to obtain body mass *m*. Wing loading was calculated as 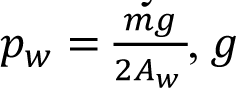 is gravitational acceleration and *_w_* is the area of one hindwing (see below). Among sampled insects, the smallest value of p_w_ was ∼1.4 Nm^-2^ in a *Brevinecroscia* sp. male, whereas the largest was ∼2300 Nm^-2^ in *Extatosoma tiaratum* females. For preliminary flight trials, high-speed videos were captured with a high-speed camera (250 fps; X-PRI, AOS Technologies AG, Switzerland; HiSpec, FasTec Imaging, USA).

Morphological data of non-phasmids were obtained from the literature. For Paleozoic insects, we used an empirical model (Wootton and Kukalová-Peck, 2000) to approximate wing loading (see **SI Datasheet 4** for details).

### 2.2 Wing shape and venation

Given that phasmids possess vestigial forewings, our sampling only evaluated hindwings. Insects were anesthetized either with a 5-min cold treatment (0 – 4 °C) or a 5-min CO_2_ treatment. We took images of fully unfolded hind wings using a digital camera (E-3, Olympus, Tokyo, Japan). Hind wing profiles were then extracted using Photoshop (Adobe Inc., USA), and were measured using ImageJ (Abràmoff et al., 2004) for wing lengths (L_w_) and wing areas (A_w_) (**Fig. 4B**). Wing aspect ratio was calculated as *AR* = *L_2_^w^*/*A_w_*.

Following Ragge (1955), we identified the anterior anal (AA) and posterior anal (PA) veins using associations with axillary sclerites at the wingbase (**SI Fig. S2**). The number of AA veins (N_AA_) with respect to p_w_ was fitted with inverse sigmoid curves using a Four Parameter Logistic Regression (R package ‘dr4pl’; Ritz et al., 2015):

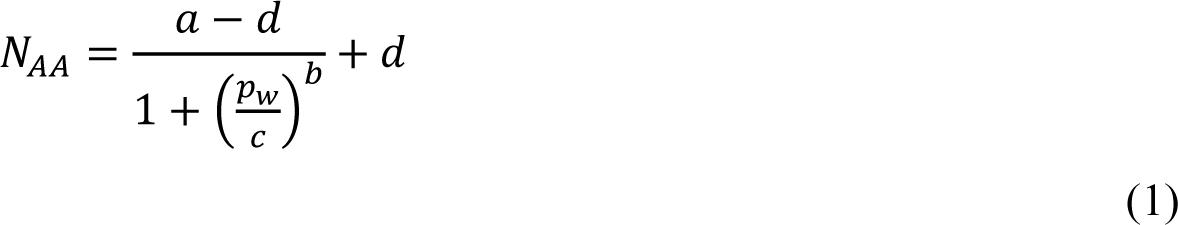

where 𝑎, 𝑏, 𝑐, and 𝑑 are the four fitting coefficients.

### 2.3 Flight muscle mass

Flight muscle mass was measured for a total of 16 different samples (6 females and 4 males reared in the laboratory, and 3 females and 3 males for each subspecies of *A. tanarata* collected in the field, with 2 – 6 individuals per sample), covering a relative wing size from 0 – 0.78 (see **SI Fig. S1**). We first cut off the metathorax from euthanized insects, and removed muscles associated with hindlegs and the median joint. We then indirectly measured the mass of flight muscles following NaOH digestion of soft tissue, with duration of chemical treatment adjusted according to the size of wings (Q > 0.5, 12 hr; 0.3 < Q < 0.5, 2 hr; Q <0.3, 30 min) (Marden, 1987).

### 2.4 Shape, area and mass of body-leg system

The dimensions and projected planforms of body-leg system were similarly collected from anesthetized insects laid dorsoventrally on a flat surface with their legs fully extended. From images, we sampled locations of wingbase (at 2^nd^ sclerite plate) and median joint (i.e., the 1^st^ abdominal tergum; Bragg, 1997), which is the axis of abdominal rotation. Given their laterally expanded abdomens and potentially different aerodynamics relative to the majority of phasmids with slender abdomens, leaf insects (Phylliidae) were excluded from this analysis.

We simplified the body-leg system as a two-link model, with the anterior body section (head + thorax) and abdomen connected at the median joint (Bragg, 1997; **SI Fig. S2**). The masses of two body sections and the legs were measured from deep-frozen specimens for 20 samples (10 female and 10 male) from 12 species (**SI Fig. S1**). Each body part was cut from frozen specimens and was measured with a portable electronic balance (PP-2060D, Acculab) with an electronic balance (R200D, Sartorius AG, Germany) indoor.

The center of mass (CM) of the body was first estimated from images of frozen specimens (with legs removed) balanced on a vertically oriented razor blade. Hind-wing mass was measured from freshly euthanized specimens. Forewings were highly reduced in size (i.e., *<* 5% A_w_ and < 0.1% mass in all sampled taxa) and were thus omitted. The total leg mass ranged from 9% – 24% of body mass (see **SI Datasheet S1**) and was included in corresponding body sections.

Based on observations of leg posture in flight (**Fig. 2**; **SI Fig. S3**), the mass of fore- and mid-legs was included in that of anterior section (𝑚_1_), and the mass of the hind-legs was included in that of the posterior section (𝑚_2_); 𝑚_1_ and 𝑚_2_ were assumed to be located at the longitudinal midpoints of the corresponding sections. The longitudinal position of the CM of body-leg system was calculated as:

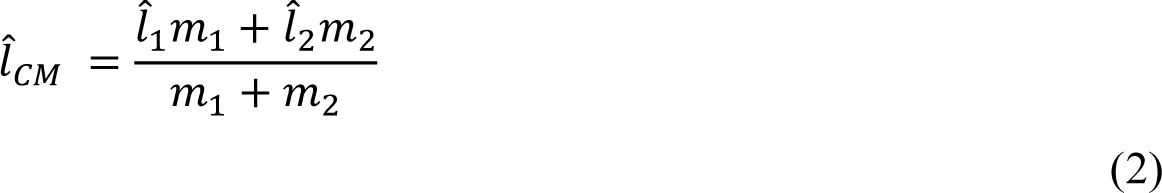

where *l̂_CM_* and *l̂_2_* are body length-normalized positions (relative to the anterior end) of 𝑚_1_ and 𝑚_2_, respectively. The radius from CM to wingbase was calculated as *r̂_wb_* = *l̂_wb_* − *l̂_CM_*, where *l̂_wb_* is the longitudinal location of the wingbase relative to the anterior end; similarly, the radius from CM to median joint was calculated as *r̂_mj_* = *l̂_mj_* − *l̂_CM_* (**Fig. 4C**).

### 2.5 Phylogenetically justified statistics

We applied log10-transformation to L_w_, L, and p_w_ in scaling analyses. We used a molecular phylogeny that includes all three subspecies of the *A. tanarata* analyzed in previous study (Zeng et al., 2020). For species lacking molecular data, we added them as polytomous tips to the node representing the latest common ancestor on the tree at the tribe or subfamily level. Each new node was added using the function ‘multi2di’ (package ‘ape’; Paradis et al. 2004), and was given a branch length randomly drawn from a normal distribution of branch lengths with a mean of 0.1 × mean branch lengths of the original tree, and a standard deviation of 0.01 × the standard deviation of branch lengths from the original tree. We then generated 100 random trees with randomly resolved polytomous tips and conducted phylogenetic generalized least square (PGLS) analyses (package ‘caper’; Orme et al., 2013) and ordinary generalized least square analyses (GLS). For each correlation, we ran PGLS on all random trees and summarized the results (ML*λ* and coefficients), which were then compared with those from GLS tests conducted without reference to the phylogeny (i.e., *λ* = 0).

For correlations of a given trait with respect to p_w_, we conducted sensitivity analyses using a series of generalized linear regressions to identify the potential p_w_ value corresponding to a critical transition in the trait. With custom-written scripts in R, we first generated a series of p_w_ ranges with different upper limits (denoted 𝑝_𝑤[𝑚𝑎𝑥]_). We then examined the significance of correlations (i.e., standard errors and *P*-values of slope coefficients) with respect to p_w_.

## 3. Results

### 3.1 Overview

Long-winged phasmids exhibited wing loadings (p_w_) of 1 – 2 Nm^-2^, which was at the same order of magnitude with those of non-phasmid insects (0.3 – 16.9 Nm^-2^; **Fig. 5**). However, sampled phasmids generally have a lower relative wing size (Q), partly due to their slender body shape, and feature a continuously increasing p_w_ with decreasing Q.

**Figure 5.**
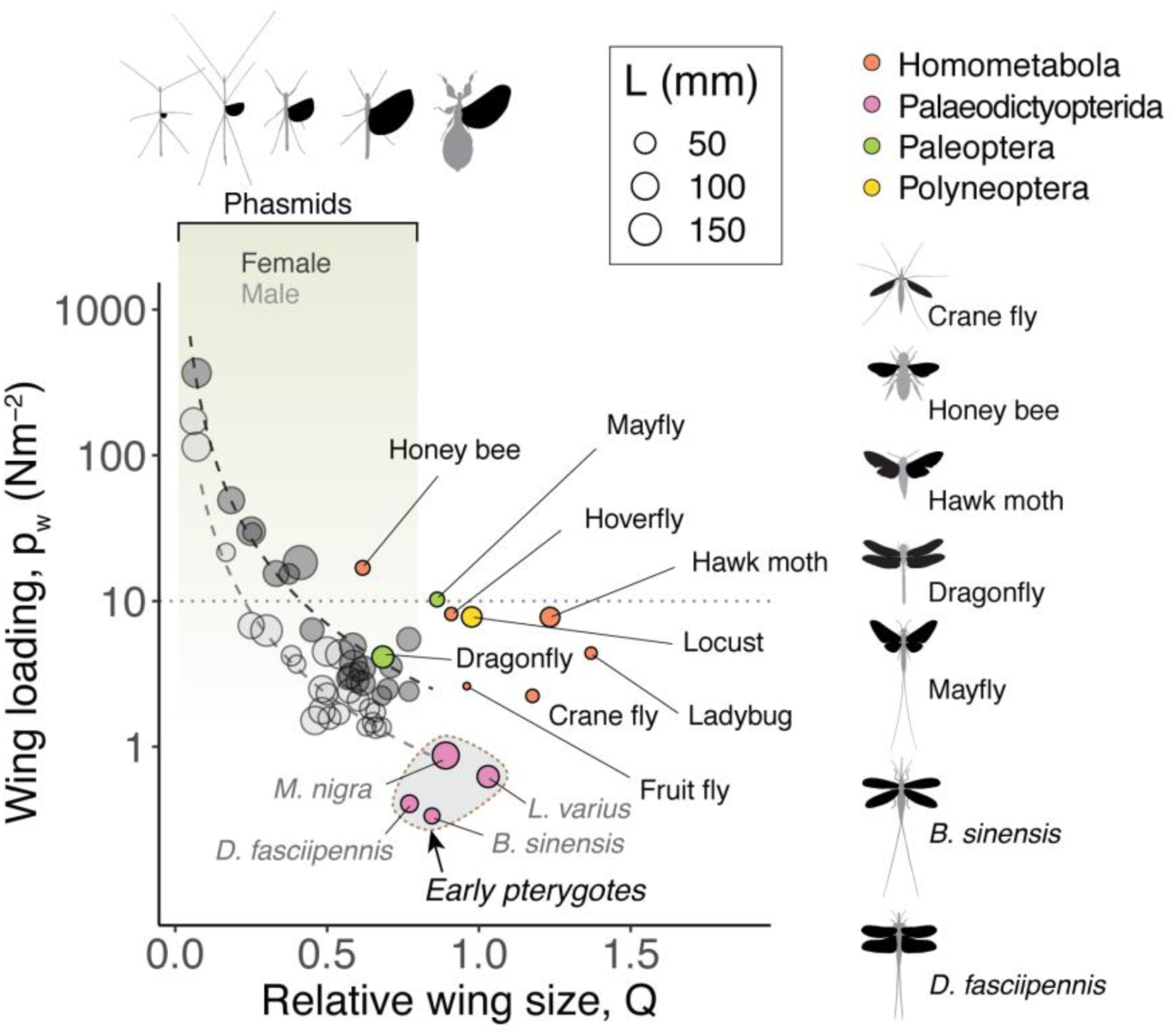
Comparisons of relative wing size, wing loading and body plan among phasmids and other winged insects. With decreasing wing size (Q), phasmids exhibit an exponential increase in wing loading (data from Zeng et al., 2020). Dot size represents body length (L). Compared with modern and Paleozoic winged insects, fully-winged phasmids have similar p_w_ but smaller Q. Morphological data for early pterygotes were extracted from the literature (*Mischoptera nigra*: Carpenter, 1951; *Lithomantis varius*: Brauckmann and Gröning, 2018; *Dunbaria fasciipennis*: Kukalová-Peck, 1971; Prokop et al., 2019; *Brodioptera sinensis*: Pecharova et al., 2015), with wing loadings estimated using a previously developed model (Wootton and Kukalová-Peck, 2000). Data for modern winged insects were obtained from numerous sources (Fruit fly, *Drosophila melanogaster*; Ladybug, *Coccinella punctata*; Crane fly, *Tipula obsoleta*; Hoverfly, *Episyrphus balteatus*; Honey bee, *Apis mellifera*; Hawk moth, *Manduca sexta*; Locust, *Schistocerca gregaria*; Dragonfly, *Aeshna juncea*; Mayfly, *Hexagenia atrocaudata*; Elltington, 1984; Heidinger, 2018; Lehmann, 1997; Stevenson, 1995; Taylor, 2003) (**SI Datasheet S4**).

Phasmid wing aspect ratio (AR) varied between 1.2 and 2.7 (male, 1.6 – 2.6; female, 1.2 – 2.7). Slender wings of AR > 2 were mostly found in males with p_w_ < 10 Nm^-2^, whereas round wings of AR < 1.5 were mostly found in partial-winged females with p_w_ > 5 Nm^-2^ (**Fig. 6**). Phasmid wings feature a unique low-AR shape, contrasting with other insect wings (either single wings or interlocked fore- and hind-wings) with an AR range of 2.5 – 6.

**Figure 6.**
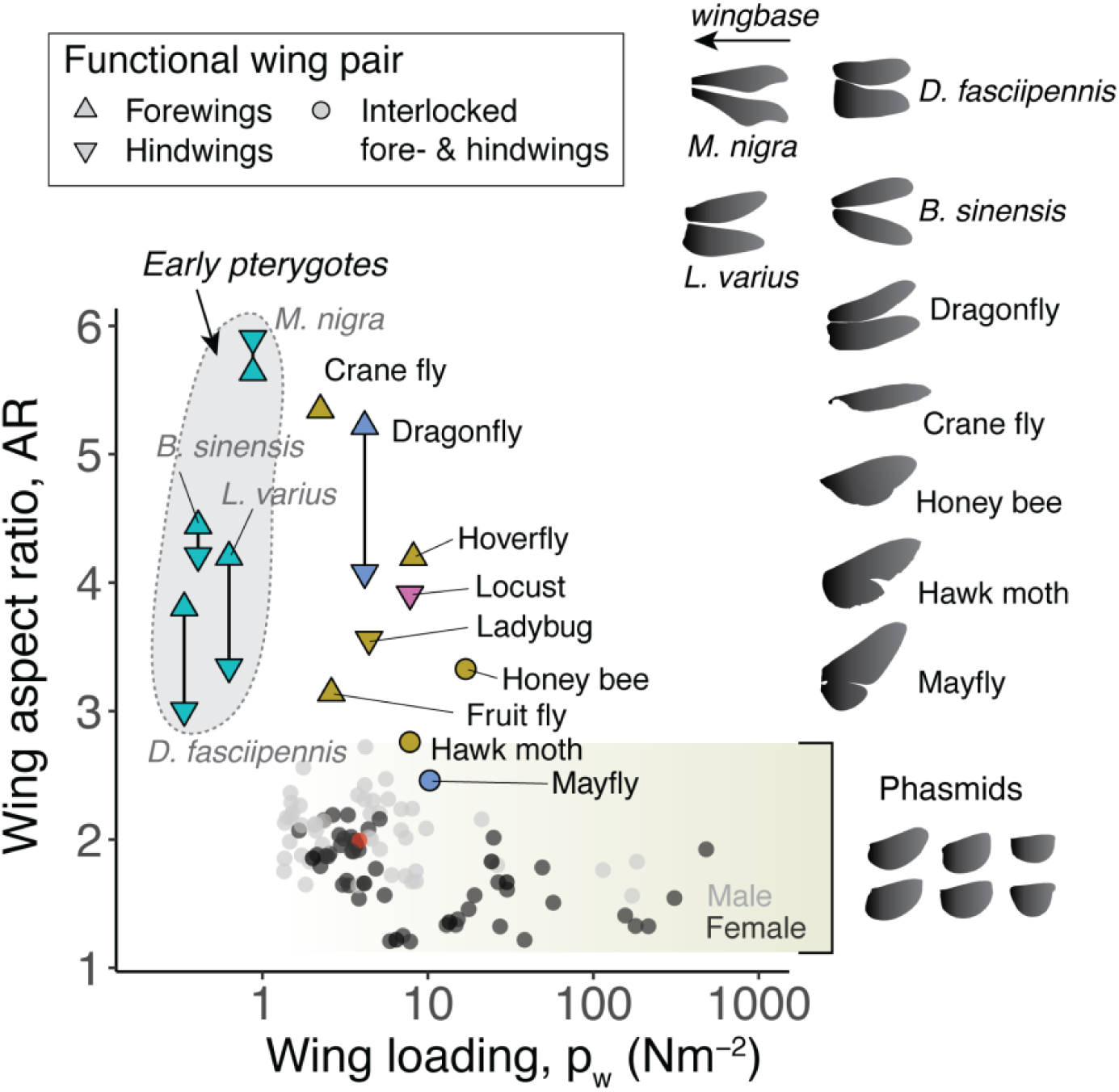
Comparison of wing aspect ratio (AR) between phasmids and other winged insects. Phasmids have rounder wings than most other taxa. For taxa with a bimotoric flight mode, fore- and hind-wings were sampled separately; otherwise, AR was calculated based on the functional wing pair (crane fly) or with two wing pairs combined as used during flight (mayfly and honeybee; see details in **SI Datasheet S4**).

### 3.2 Phasmid wing shape scales with wing size

Phasmid wing AR was not correlated with body size (L), but was significant correlated with wing size-derived principal variables (**SI Datasheet S2**). On one hand, wing AR of male phasmids was positively correlated with the absolute wing length L_w_:

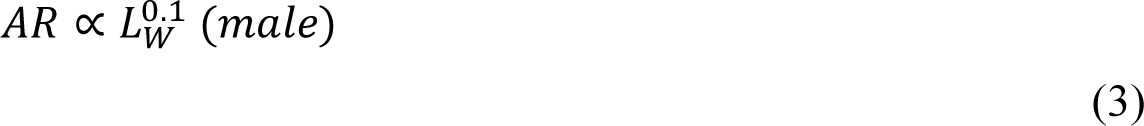

where the scaling exponent 0.1 suggests mild expansion of the mean wing chord size (𝐶_𝑤_) as the wings become longer (**Fig. 7A**). As *AR* = *_w_C_w_^−1^*, we can rewrite Eqn. (3) as *L_w_C_w_*^−1^ ∝ *L*_*w*_^0.1^, leading to a 𝑤 𝑤 𝑤 𝑤 𝑤 negative allometry of mean chord size *C_w_* ∝ *L*_*w*_^0.9^.

**Figure 7.**
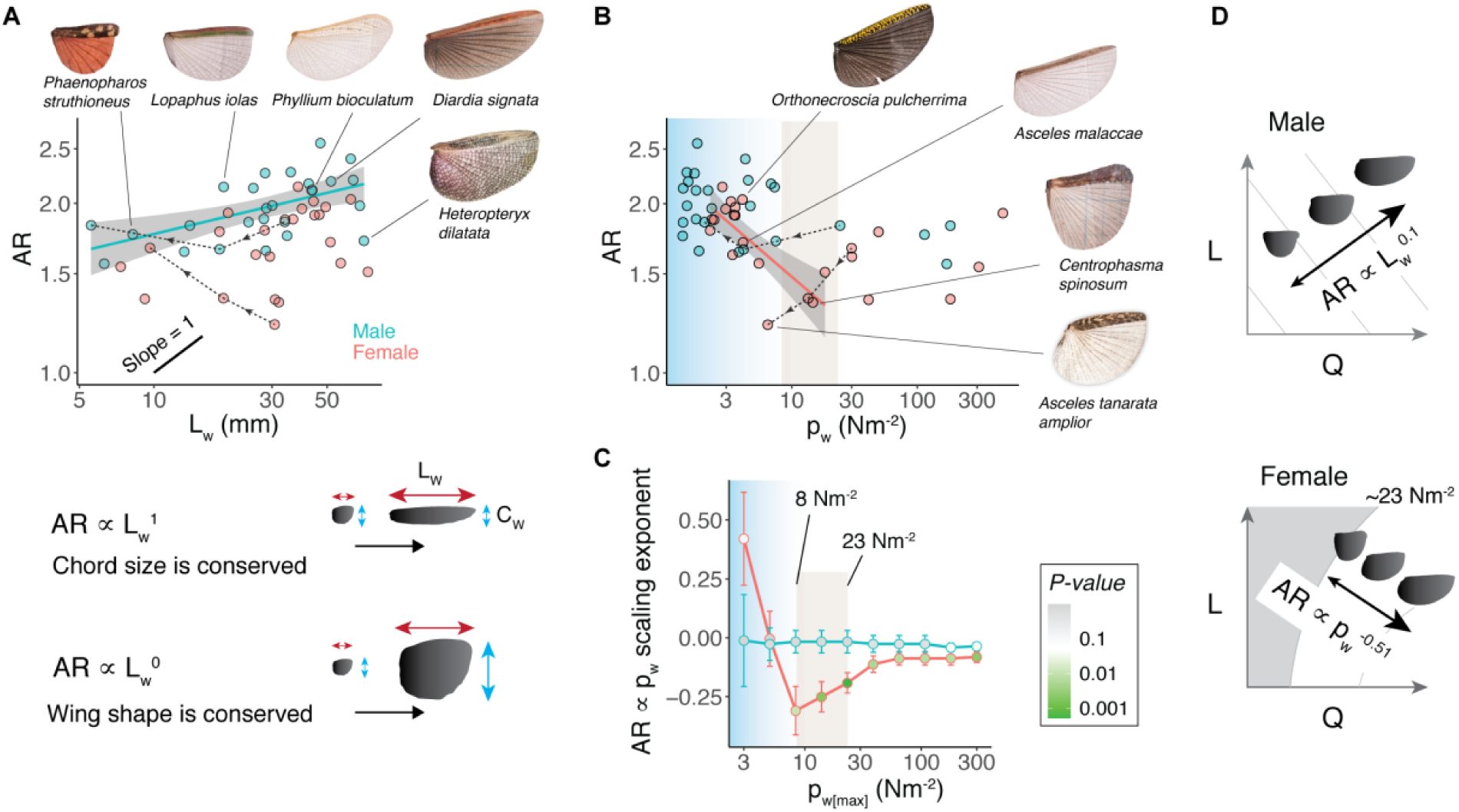
Sex-specific scaling of phasmid wing shape. **(A)** Significant scaling of wing aspect ratio (AR) with wing length (L_w_) was found in males (slope of 0.10±0.03; *P* < 0.01) but not in females (*P* = 0.12). This result indicates an elongation with chordwise expansion, as explained with two reference models at the bottom. **(B)** Wing AR scales with wing loading (p_w_) in female phasmids. A clear reduction of AR in female phasmids was observed in p_w_ range of 4 – 30 Nm^-2^, whereas wings of male phasmids showed no such reduction. Such trends are further elucidated with a sensitivity analysis in **(C)**, wherein generalized linear regression models were performed with different upper limit of wing loading, 𝑝_𝑤[𝑚𝑎𝑥]_. The strongest correlation corresponded with 𝑝_𝑤[𝑚𝑎𝑥]_ = 10^1.4^ Nm^-2^, as shown by the trend line (slope of -0.51±0.14, *P* < 0.01). **(C)** The variation of the scaling exponent as plotted against p_w_, with error bars representing one standard error, and colors representing the *P*-value. For females, the lowest slope coefficient corresponds with 𝑝_𝑤[𝑚𝑎𝑥]_ = 10^0.92^ (≈ 8) Nm^-2^, whereas the lowest *P*-value corresponds to 𝑝_𝑤[𝑚𝑎𝑥]_ = 10^1.36^ (≈ 23) Nm^-2^, as shown by the trend line in **(B)**. For males, the scaling was non-significant and insensitive to the choice of 𝑝_𝑤[𝑚𝑎𝑥]_. **(D)** Summaries of trends for wing shape variation in male (top) and female (bottom) phasmids with respect to L and Q. In **(A)**-**(B)**, arrows with dashed lines represent variation within *A. tanarata* subspecies along increasing altitude (see **SI Section A**).

On the other hand, wing AR of female phasmids exhibited a reduction with increasing p_w_, as shown by a progressively rounder wing shape (**Fig. 7B**). Using sensitivity analysis (**Fig. 7C**), we identified that p_w_ ≈ 23 Nm^-2^ was the upper limit of p_w_ that corresponded with the most significant AR-p_w_ correlation.

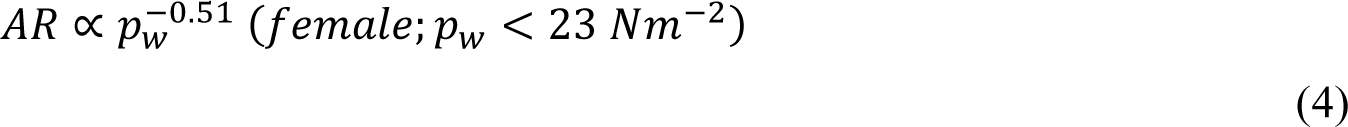

The shapes of partial wings thus follow linear transitions with sex-specific trends (summarized in **Fig. 7D**), which likely reflect secondary aerodynamic adaptation along with influences from body mass allometry (see **Discussion**).

### 3.2 Scaling of wing venation

The first observed pattern was that larger wings possess more radial veins, but the two vein groups showed different trends of variation (**Fig. 8A,B**). Although correlated with absolute wing size L_w_ (scaling exponent ∼0.37), the number of radial veins (𝑁_𝑣_) was most significantly correlated with the relative wing size Q and wing loading p_w_ (**SI Datasheet S2**). Examining more closely the two radial vein groups, we found that the number of anterior anal veins (𝑁_𝐴𝐴_) only varied from 5 – 7, whereas that of the posterior anal (PA) veins (𝑁_𝑃𝐴_) was more variable (between 2 – 14). With increasing p_w_, 𝑁_𝑃𝐴_ exhibited a linear reduction, whereas 𝑁_𝐴𝐴_ showed a sharp reduction from 7 to 5 near p_w_ ≈ 10 Nm^-2^ (**Fig. 8C**), with the value of 𝑁_𝐴𝐴_ = 6 characterizing a critical transition. Using logistic regression models (**Eqn. 1**), we found the sharp reduction of 𝑁_𝑣_ corresponded with a critical pw ≈ 10 Nm^-2^ in both sexes (10^0.8^ Nm^-2^ for males and 10^1.3^ Nm^-2^ for females) (**Fig. 8D,E**).

**Figure 8.**
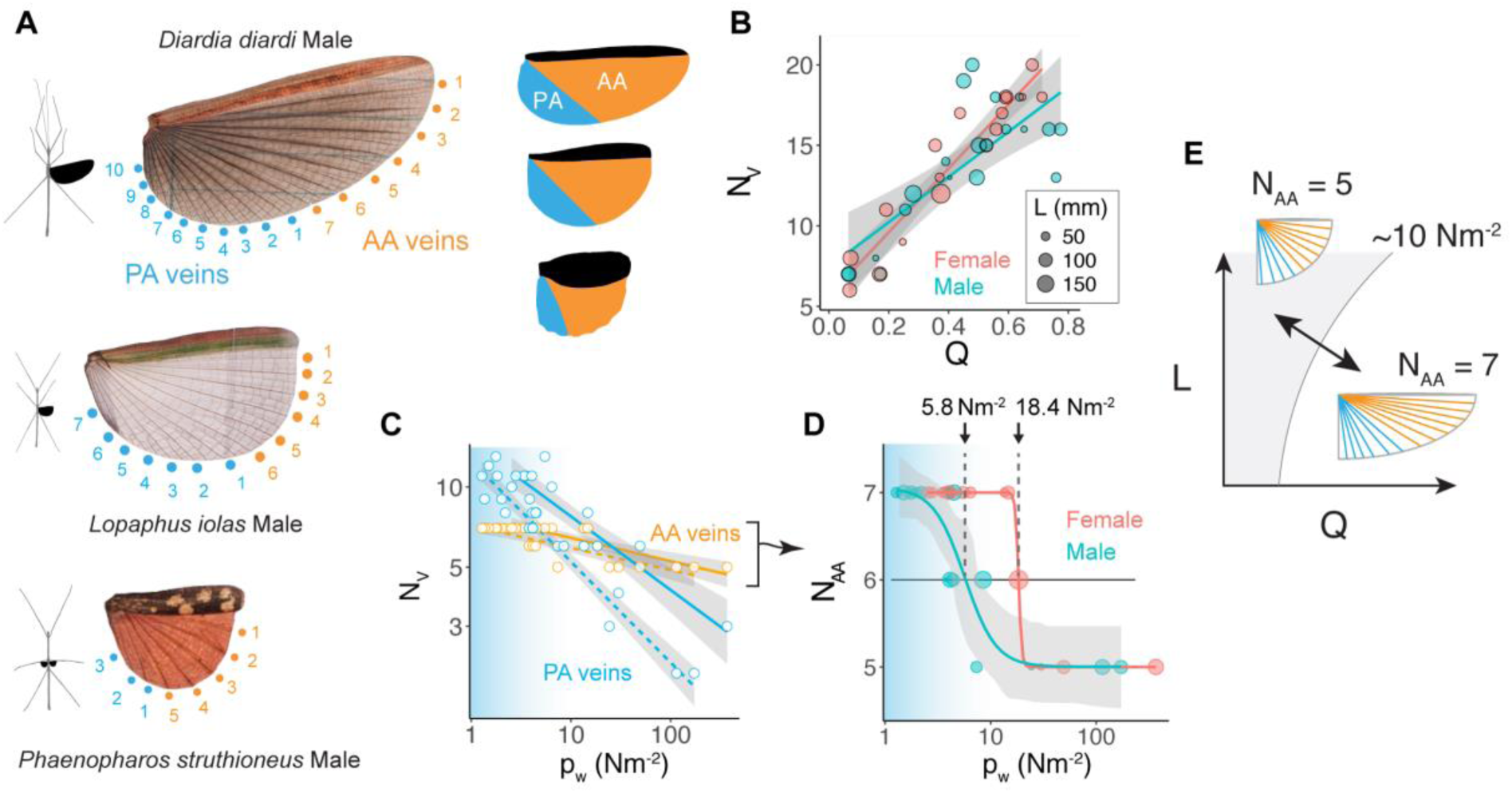
The number of primary radial veins scales with wing loading. **(A)** Exemplar wings showing reduction of primary radial veins. (Right) Different area proportions for two vein groups: (1) the relatively conserved anterior area supported by AA veins (orange), and (2) the wing size-correlated coverage by PA veins (blue). **(B)** Correlation between the total number of radial veins (N_V_) and the relative wing size Q (slope: females, 19.8±1.7; males, 14.0±2.7; means±s.e.; *P* < 0.0001 for both sexes). Relative dot size indicates body length. **(C)** The number of PA veins varied between 2 – 14 (blue), and is linearly correlated with p_w_ (females: -0.39±0.04; males: -0.37±0.02), whereas the number of AA veins (N_AA_) is more conserved, varying from 5 – 7 (orange). Trend lines are based on linear regression models. **(D)** The sharp reduction of N_AA_ from 7 to 5 across a critical p_w_, as fitted with sigmoid functions (**SI Datasheet S1**). The model-predicted critical p_w_ at N_AA_ of 6 is 10^0.8^ Nm^-2^ (or ∼5.8 Nm^-2^; confidence interval, 4.4 – 8.1 Nm^-2^) for wings of males, and 10^1.3^ (or ∼18.4 Nm^-2^; confidence interval ∼0) for wings of females. **(E)** Schematic summary for the sharp reduction of the number of primary radial veins across a critical p_w_ ≈ 10 Nm^-2^.

The second general pattern we noticed was that long wings had more radial vein elements, specifically the two levels of intercalary veins (namely primary and secondary) found along the membrane margin; short wings lacked the one or both levels of intercalary veins (**Fig. 9A**). A preliminary explanation of this pattern is that, as the inter-vein angles are relatively conserved (e.g., 6.8° – 8.8°; see **SI Fig. S6D**), gap size between adjacent veins should increase toward the wing margin, with greater separation in longer wings; the presence of intercalary veins (and cross veins) would help reinforce both transverse stiffness and fracture toughness along the wing margin.

**Figure 9.**
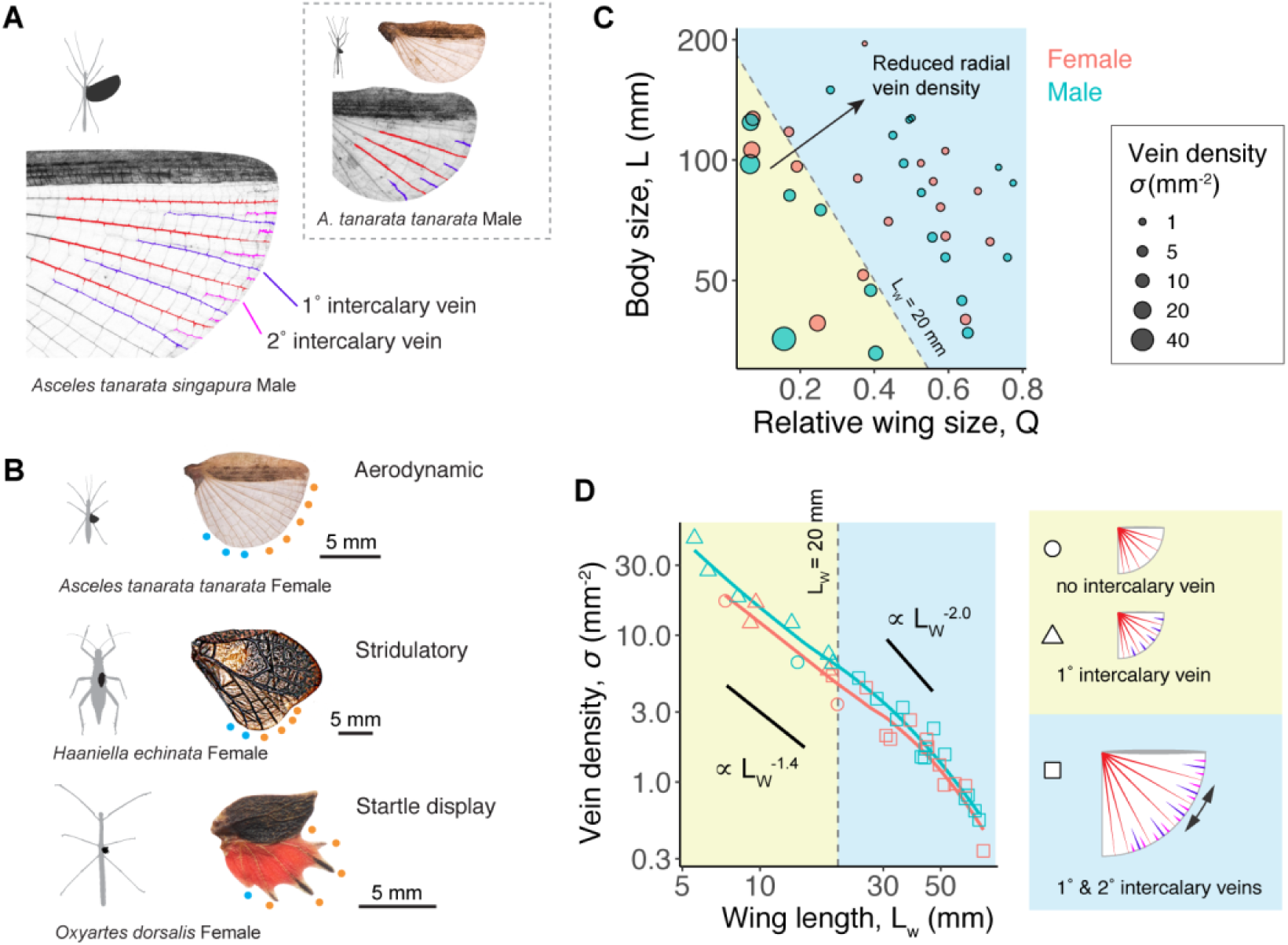
Secondary venation changes with wing size. **(A)** As gap size between adjacent radial veins increases toward the wing margin, intercalary veins are developed. Long wings possess two levels of intercalary veins, whereas short wings possess only one, as demonstrated here using *A. tanarata singapura* and its high-altitude, short-winged relative *A. tanarata tanarata*. **(B)** Examples of short wings of length <20 mm. The top wing with a complete membrane potentially serves rudimentary aerodynamic function, contrasting with two other wings serving non-aerodynamic functions. **(C)** Mapping of radial vein density (𝜎 = N_V_/ A_w_, as represented by dot size) with respect to L and Q, showing a decreasing trend as wing size L_w_ (=LQ) increases. **(D)** The power-law scaling of 𝜎 with L_w_. Point shapes represent the type of venation. A critical wing size of L_w_ ≈ 20 mm separates two regimes of secondary venation, as highlighted by the background coloration. (Right) Schematic demonstrations of two regimes: (1), in long wings (L_w_ > 20 mm), two levels of intercalary veins fill the gaps between primary radial veins near wing margin; and (2), in short wings (L_w_ ≤ 20 mm), there is either one level of intercalary veins or none.

To further explain how such a pattern is correlated with the distribution of radial veins on differently sized wings, we first show that vein density (𝜎 = 𝑁_𝑣_/𝐴_𝑤_) decreased with increasing wing length (**Fig. 9A**), which is in accordance with aforementioned negative allometry of N_v_ with L_w_ (𝑁_𝑣_ ∝ 𝐿^0.37^). Next, such a negative scaling is more clearly seen by plotting 𝜎 against L_w_ (**Fig. 9B**), such that two levels of intercalary veins were only found in long wings with L_w_ > 20 mm. Correspondingly, the scaling of vein density differed between long and short wings across L_w_ = 20 mm:

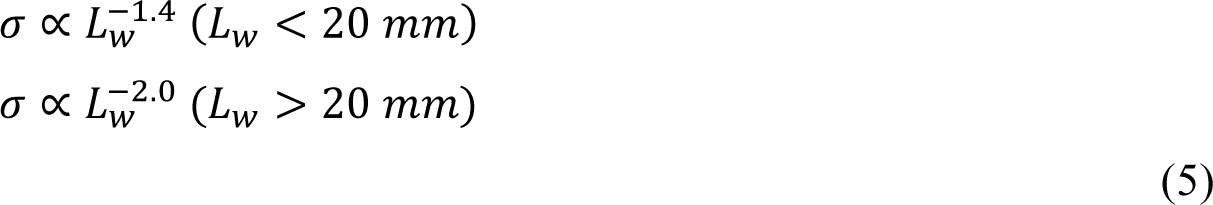

The lower scaling exponent for longer wings corresponds with the increase of wing area in spanwise direction as wing size increases (i.e., increasing AR) (**Fig. 7A**).

### 3.3. Scaling of wing and flight musculature masses

First, over a wing length range 0.5 – 7.7 cm, the mass of a single wing (𝑚_𝑤_) varied between 0.4 mg and 160 mg, which is heavier than other insect wings of similar lengths (0.2 – 36 mg) (**Fig. 10A**). Whereas wings longer than 5 cm generally weighed more than 0.1 g in both sexes, the wings of males exhibited a steeper slope of mass with respect to wing size:

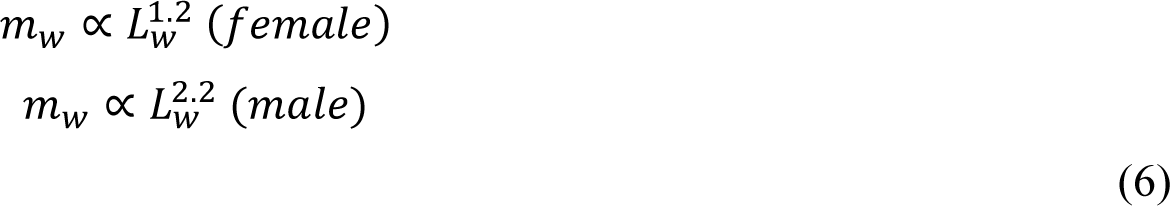

**Figure 10.**
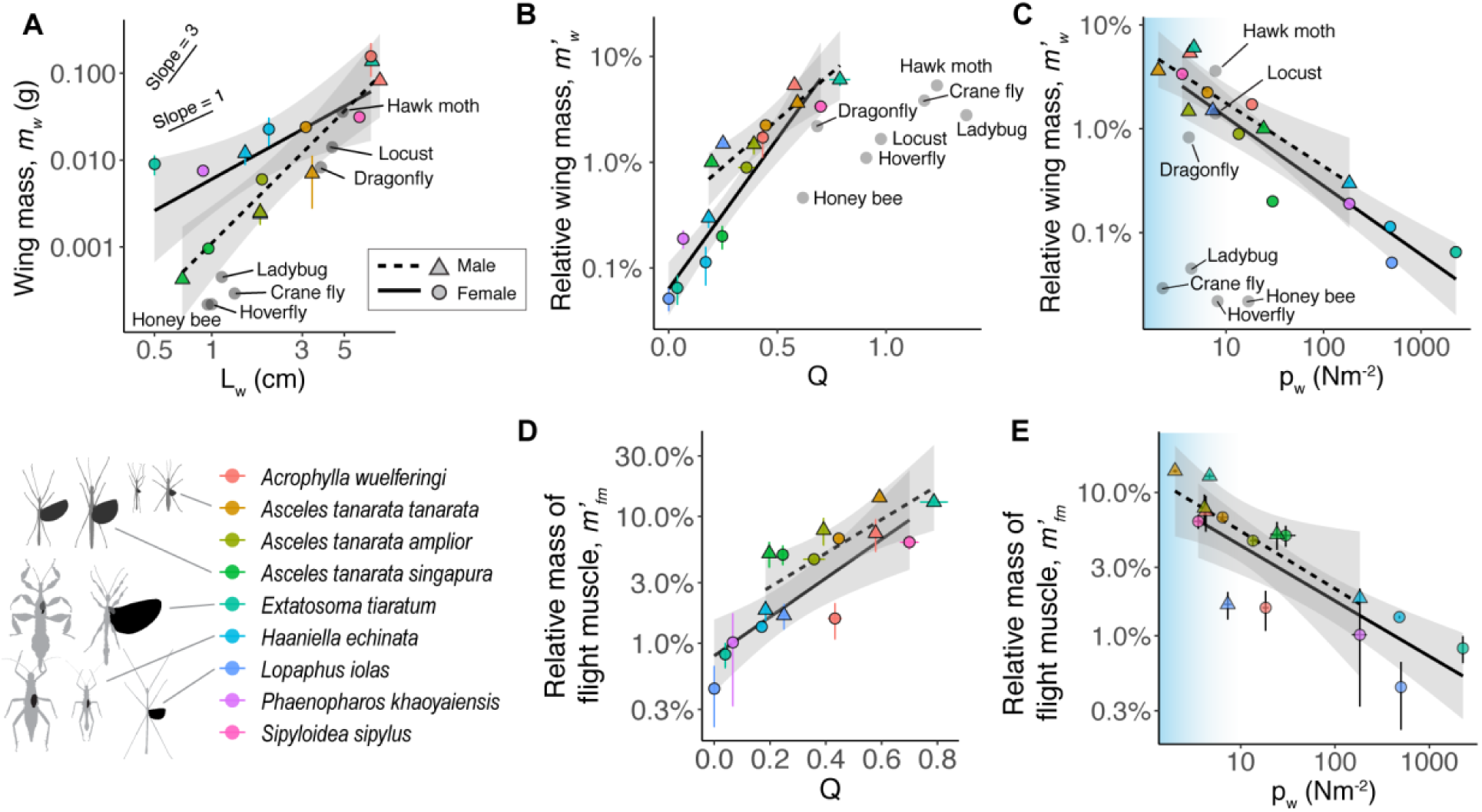
Scaling of wing mass and flight muscle mass. **(A)** Scaling of the mass of a single hindwing with respect to wing length (L_w_) (slope: females, 1.2±0.4; males, 2.2±0.4; mean±s.e.). **(B)** The scaling of relative wing mass (for both hindwings combined) with respect to relative wing size Q, showing that females possess relatively lighter wings than males (slope: females, 0.048±0.007; males, 0.092±0.014). Compared to other winged insects, stick insects possess relatively shorter but also heavier wings. **(C)** The scaling of relative wing mass with wing loading (slope: females, -0.010±0.003; males, -0.608±0.154). **(D)** The scaling of relative mass of flight muscle with respect to relative wing size, showing that males have greater volume of flight muscle than females (slope: females, 0.086±0.025; males, 0.185±0.046). **(E)** The scaling of relative mass of flight muscle with wing loading (slope: females, -0.386±0.085; males, -0.402±0.179). All values represent sample-specific means±S.D.; dot shape and line type represent different sexes, colors represent different species, and trend lines are based on linear regression models.

If wings are simplified as plates with uniform thickness and constant planform shape, these power-law scaling relationships suggest a moderate increase of the mean thickness with increasing 𝐿_𝑤_; this allometry is intermediate to two models: (i) wing thickness is conserved and mass is proportional to wing length (exponent of 1), and (ii) wing dimensions increase isometrically in all three dimensions (exponent of 3).

Second, at the whole-insect level, the relative mass of phasmid wings (two hindwings combined; 𝑚^′^ = 2𝑚_𝑤_/𝑚) exhibited linear scaling with Q and pw. Long-winged phasmids had similar 𝑚^′^ with other insect wings, and the highest was 6% found in male *Extatosoma tiaratum* (Q ∼0.8). With decreasing Q, 𝑚^′^ continuously reduced and reached < 1% in miniaturized wings (**Fig. 10B**). With respect to increasing 𝑝_𝑤_, 𝑚_𝑤_′ showed an inverse scaling (**Fig. 10C**), showing that the evolution of lower p_w_ (e.g., smaller body size or longer wings) leads to relatively heavier wings (see **Discussion**).

Third, the relative mass of flight muscle (𝑚^′^) was mostly below the minimum for takeoff described in other volant insects (12% – 16%; Marden, 1987); with reducing Q, 𝑚^′^ continuously dropped from 14% to ∼0% (**Fig. 10D**). Similar to wing mass, 𝑚^′^was correlated with Q and pw (**Fig. 10E**). Neither wing mass and flight muscle mass were correlated with body size (**SI Datasheet S2**).

### 3.4 Scaling of shape and area of body-leg system

Among sampled phasmids with slender body shape, the body aspect ratio (i.e., the ratio of length to mean diameter; 𝐴𝑅_𝑏_) varied from 10 – 56 (15 – 56 in males, and 10 – 25 in females), and was positively correlated with body length 𝐿 in males (𝐴𝑅_𝑏_ ∝ 𝐿^2.82^), but not in females (**Fig. 11A**). This high range of slenderness was much larger than that in all other winged insects used for comparison (mostly <10, except for a dragonfly).

**Figure 11.**
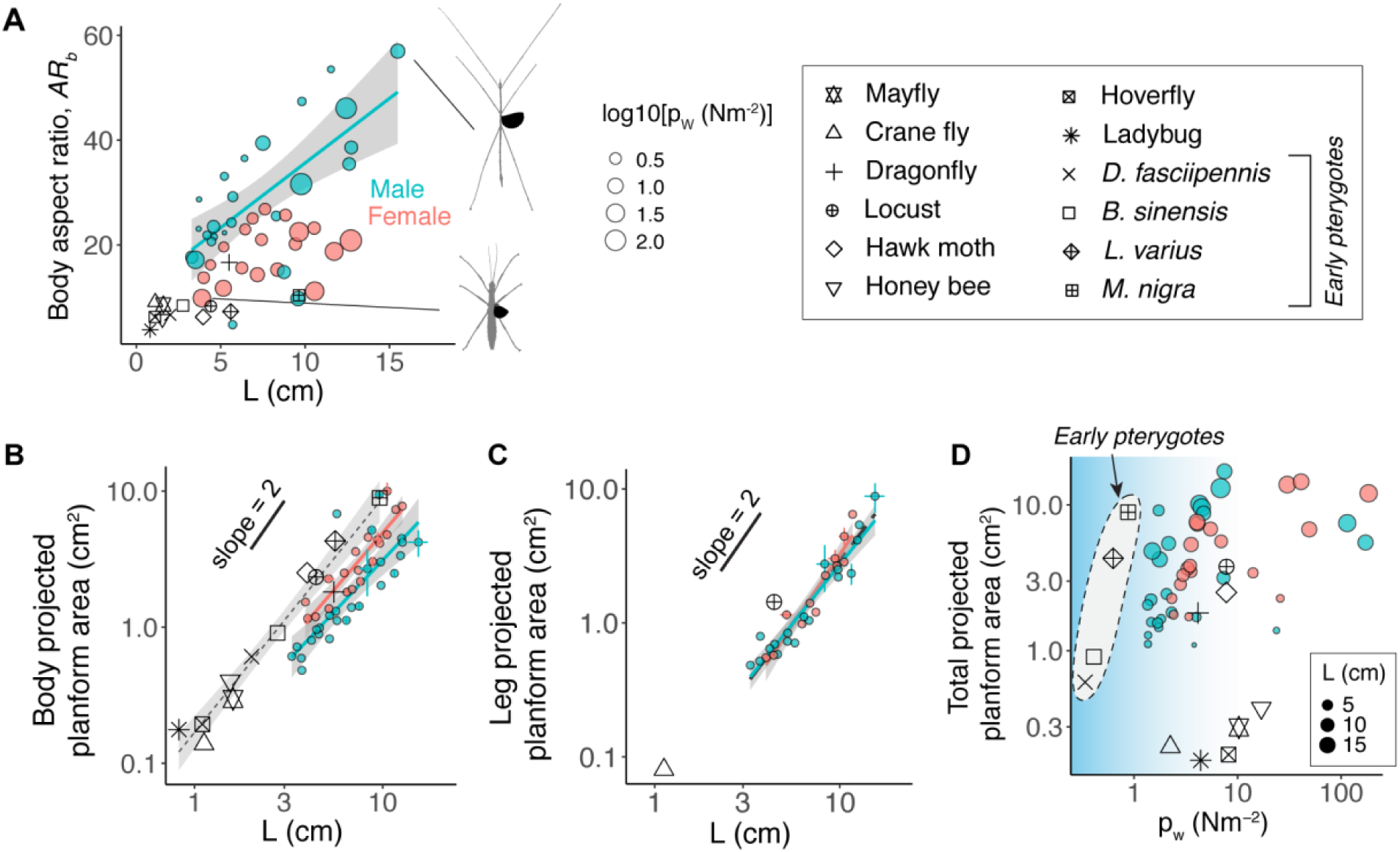
Phasmid body-leg systems feature diverse shapes and relatively greater projected areas. **(A)** Body aspect ratio (𝐴𝑅_𝑏_) versus body length (L) in phasmids and other winged insects. A significant correlation between 𝐴𝑅_𝑏_ and L was found for males (slope of 2.82±0.38; mean±s.e.; *P* < 0.0001), but not for females (*P* > 0.05). Point size represents p_w_, which was not correlated with 𝐴𝑅_𝑏_. **(B)** The projected planform area of the body scales with L. The greater slope intercept in females corresponds to wider body segments (intercept: females, -0.97±0.19; males, -0.96±0.07; sex-averaged, -1.00±0.15; slope: females, 1.65±0.22; males, 1.35±0.09; sex-averaged, 1.54±0.15; *P* < 0.0001 for all correlations). Dotted trend line represents the scaling for other insects (intercept: -0.77±0.06; slope, 1.72±0.12; *P* < 0.0001). **(C)** The projected planform area of all legs scales with L, with a sex-averaged intercept of -1.32±0.11 and slope of 1.74±0.13 (*P* < 0.0001). Data were available from two other insects (crane fly and locust) for comparison. **(D)** The dorsally projected area of the body-leg system during flight plotted against wing loading p_w_, whereby phasmids generally have greater aerodynamic surface area than other winged insects of similar p_w_. Dot size represents phasmid body length. For non-phasmid insects, the total projected area consists of dorsally projected body area if legs are folded during flight (see details **SI Datasheet S4**).

The projected planform areas of the body and legs both ranged between 0.5 – 10 cm^2^ and were not correlated with either Q or p_w_. Instead, they mainly scaled with body length 𝐿, with the power-law exponents averaging 1.54 for body area and 1.74 for area of legs (**Fig. 11B,C**). Scaling exponents <2 for body and leg area correspond with disproportionate increases in slenderness as body-leg segments become longer. Due to large area of both body ang legs, phasmids possessed much greater aerodynamic surface area than taxa with similar wing loading (**Fig. 11D**).

### 3.5 Conservation of mass distribution

Despite the wide range of p_w_ (female, 3.3 – 500 Nm^-2^; male, 1.5 – 183.6 Nm^-2^), the relative mass of the anterior body sections (head + thorax) was similar between the two sexes (female, 37.5±2.9%; male, 39.9±5.7%; mean±S.D.), but the abdomens of female phasmids were heavier by ∼16% body mass relative to those of males (female, 49.4±3.8%; male, 33.3±3.9%) (**Fig. 12A**). Contrasting to the relatively conserved abdomen mass in female phasmids, a significant reduction of abdomen mass with increasing p_w_ was found in male phasmids (**Fig. 12B**). Also, male phasmids had relatively heavier legs than females (relative leg mass: female, 11.4±2.1%; male, 22.5±7.0%), with ∼60% of the mass from the combination of fore- and mid-legs (female, 57.9%; male, 59.2%) (See **SI Datasheet S1**).

**Figure 12.**
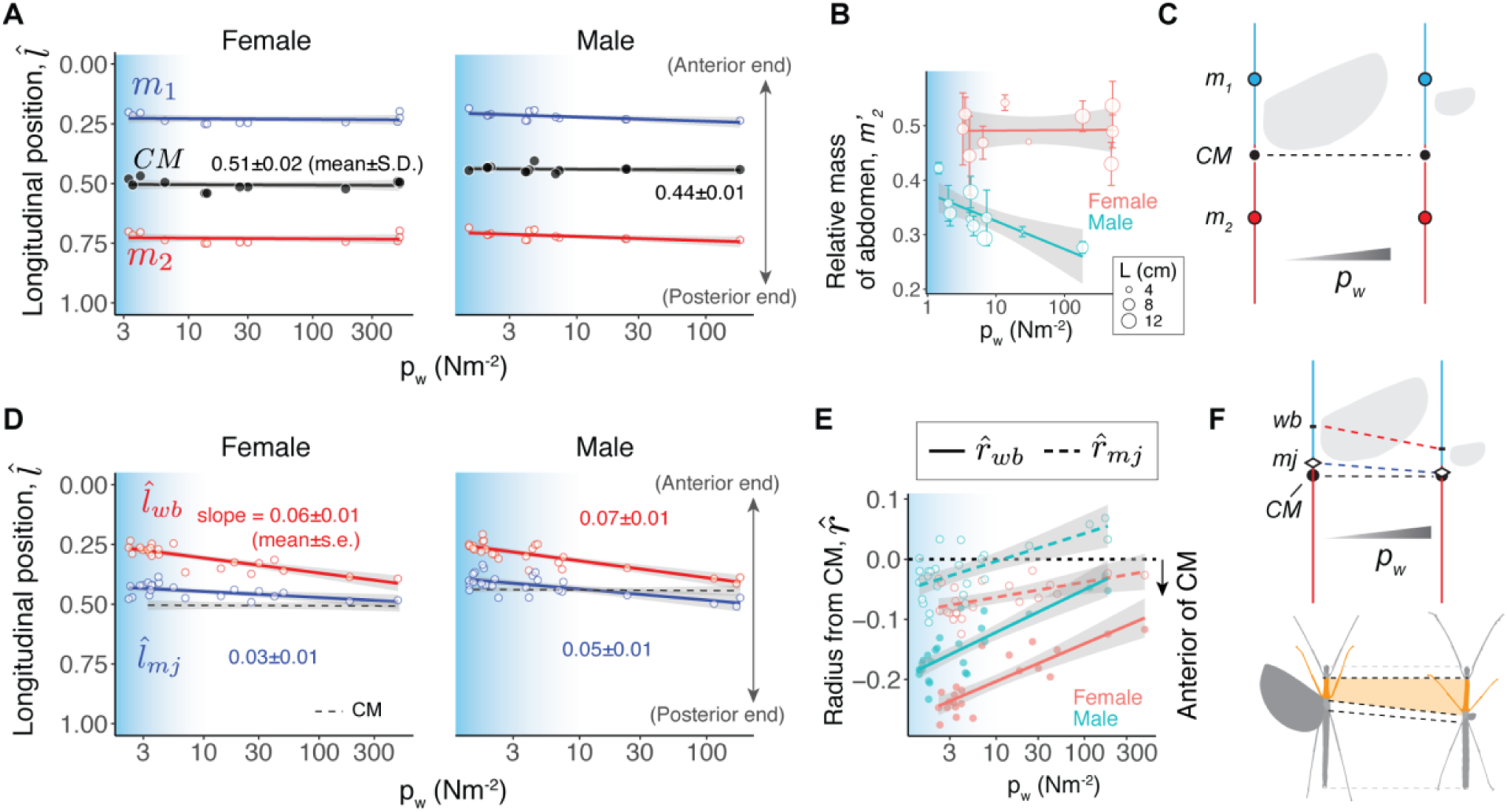
Mass distribution and body plan in phasmid flight transition. **(A)** Longitudinal locations of body-leg section masses plotted against wing loading (p_w_). Blue and red dots represent midpoints of the anterior and posterior sections, respectively (Fig. 4C); black dots represent the body-leg system center of mass (CM). Trend lines are based on linear regression models. The location of CM (*l̂_CM_*) is not correlated with p_w_ (*P* > 0.05). **(B)** The relative mass of abdomen decreases with increasing p_w_ in males (slope of -0.05±0.02, mean±s.e., *P* < 0.05), but not in females (*P* = 0.96). **(C)** Schematic illustration of the conserved location of CM across a wide range of p_w_. **(D,E)** Longitudinal locations of wingbase (*l̂_wb_* , red) and median joint (*l̂_mj_* , blue) shift posteriorly with increasing p_w_, changing the radii from CM to wingbase (*r̂_CM_*) and median joint (*r̂_CM_*). Specifically, *r̂_CM_* shifts posteriorly with increasing p_w_ (slope: females, 0.04±0.02; males, -0.05±0.02; *P* < 0.05 for both); 𝑟_𝑚 𝑗_ was generally <10% body length (intercept, male, -0.05±0.01; female, -0.09±0.01, *P* < 0.001; slope, male, 0.05±0.01; female, 0.03±0.01, *P* < 0.01). Note the y-axes are flipped in **(A)** and **(D)** to match the insects’ orientation in schematic drawings. **(F)** A schematic demonstration for the shifts in wingbase (wb) and median joint (mj) with respect to wing reduction, which is coupled with elongation of the mesothoracic segment (orange), as demonstrated with two exemplar profiles (left, *Asceles malaccae* female; right, *Phaenopharos struthioneus* female). See **SI Datasheet S1** for details.

Combining the relatively masses of body and legs, we estimated the longitudinal location of body-leg center of mass (*l̂_CM_*), and found it was highly conserved in both sexes (female, 0.51±0.02; male, 0.44±0.01) (**Fig. 12A,C**). In general, except for the abdomen mass in males, all other relative masses were not correlated with L, Q or p_w_ (see **SI Datasheet S2**). At the species level, however, an increase in the relative mass of legs was correlated with body miniaturization in *A. tanarata*, which might be a specific case of developmental tradeoffs (see **SI Section A**).

### 3.6 Shift of pterothoracic segment location

In phasmids, the main pterothoracic segment (metathorax) bears the functional wing pair, and its posterior end is connected with the median joint, which is relevant to abdominal movement. In long-winged phasmids (p_w_ < 3 Nm^-2^), the longitudinal position of wingbase (*l̂_CM_*) was ∼0.26 (female, 0.26±0.02, mean±S.D.; male, 0.26±0.03), which falls in the range of 0.12 – 0.26 reported for other volant insects (Ellington, 1984). In short-winged phasmids (p_w_ > 100 Nm^-2^), however, the wingbase shifted posteriorly, with *l̂_CM_* increased to ∼0.40 in both sexes (female, 0.39±0.01; male, 0.40±0.01) (**Fig. 12D**).

Correspondingly, the radius from CM to wingbase (*r̂_mj_*) was reduced by ∼15% body length in both sexes (**Fig. 12E**). A similar shift was also found with the median joint, the longitudinal position (^𝑙^^) of which shifted by ∼0.05 in females (from 0.44±0.03 in long-winged females to 0.49±0.00 in short-winged females) and by ∼0.09 in males (from 0.40±0.04 to 0.49±0.02) (**Fig. 12D**). The radius from CM to median joint (𝑟_𝑚𝑗_) was generally <10% body length and also shifted posteriorly with increasing p_w_, even becoming posterior of CM in short-winged males (**Fig. 12E**). As both landmarks are associated with the metathorax, a longitudinal shortening of the metathoracic segment with increasing p_w_ was also evident given the greater reduction of *l̂_wb_* relative to *l̂_mj_* . Such a reconfiguration is also associated with a relative elongation of the mesothoracic segment and longitudinal compression of the abdominal region (**Fig. 12F**).

## 4. Discussion

We showed that flight-related morphological traits evolve following linear scaling relationships, but with specific principal variables often suggesting influences from both developmental regulation and flight-related selection. Below, we first address (1) what we can learn about flight in fully-winged phasmids and (2) what we can then learn about flight transition in phasmids (**Fig. 1C**). Lastly, we discuss what we can infer about insect flight transitions in general, especially in early pterygotes (**Fig. 1B**).

### 4.1 Characteristics of phasmid flight apparatus

Fully-winged phasmids may be used as a model for (1) comparing powered flight of phasmids and with that of other insects, and (2) comparing flight performance among phasmids with different sized wings. Compared to other winged insects, phasmids exhibit a series of unique features in wing design and body-leg system. Wings of slender-bodied phasmids feature narrower, cambered and elytrized costal edges that cover their slender abdomens. This apparently contributes to their visual mimicry as twigs or visual camouflage when resting on vegetational structures (**Fig. 3A**). By contrast, a less condensed costal edge can be seen in wings of leaf phasmids, which have wide, flattened abdomens (**SI Fig. S2**). The wing membrane supported by primary radial veins is also unique because it can be folded in a fanlike fashion, and undergoes torsional and flexural deformations during stroke cycles (**Fig. 2; SI Fig. S3**). The intercalary veins and associated cross-veins distributed along the wing margin likely strengthen transverse tensile stiffness and enhance fracture toughness (Dirks and Taylor, 2012).

Phasmid wings are unique in terms of highly variable sizes and shapes derived from a conserved template mostly composed of two components: an elytrized costal edge, and a flexible membrane. This design is simplified and modularized with repeating vein components compared to other insect wings, even those of orthopteroid insects (e.g., mantis and locust; Smart, 1953; Herbert et al., 2000). Little is known about the development of phasmid wings, but this design may partly explain why wing size is so evolutionarily plastic in phasmids, given that regulated duplication of radial vein components is developmentally coupled with wing size (**Fig. 8**).

One pending question is the wing aerodynamics in long-winged phasmids (capable of ascending flight), such that there are potential tradeoffs between wing-folding mechanisms, aerodynamic efficiency, and wing shape. Fanlike-folding may jeopardize flexural stiffness (especially in the chordwise direction), whereas increased wing area (and thus lowered aspect ratio; **Fig. 7B**) may be an adaption for enhancing aerodynamic efficiency. The costal edge should be narrow enough to overlap the abdomen and at the same time remain thick to maintain sufficiently high spanwise stiffness (Combes and Daniel, 2003), which likely cause the wings to be heavier than other insect wings of similar size (**Fig. 10A**).

Whereas our data on wing loading (**Fig. 5**), relative mass of flight muscle (**Fig. 10D**) and flight behavior (**Fig. 2**) suggest fully-winged phasmids are good flyers, their flight is likely to be energetically costly due to large drag area of the body-leg system (**Fig. 11D**). Given their slender bodies and legs, aerodynamic and inertial forces result in relatively long moment arms, and thus their stability is sensitive to leg and abdomen postures and movements. Detailed studies of the wing and body-leg kinematics for free-flying phasmids will help clarify these issues.

### 4.2 Transitions in wing morphology

Partial wings of phasmids are evolutionary consequences of diverse selective regimes (Zeng et al., 2020). As summarized in **Fig. 13A**, wing-related parameters are correlated with both relative wing size Q and wing loading p_w_, implying both developmental truncation and secondary flight-related selection. The main evidence for developmental truncation would be structural reduction and simplification (Hanken and

**Figure 13.**
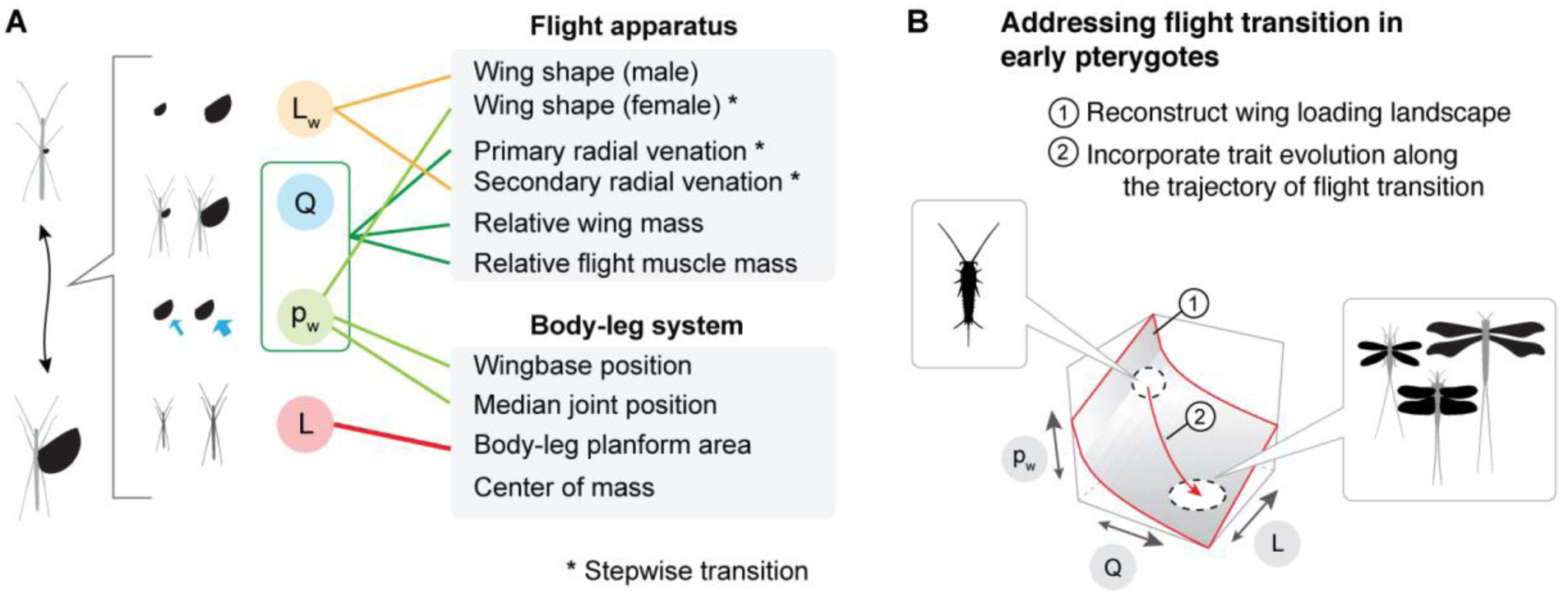
Morphological scaling in phasmid flight transition, and implications for flight gain in early pterygotes. **(A)** Summary of scaling relationships. Significant correlations are represented by lines connecting the principal study variables (left) and other morphological features (right). **(B)** Future steps for evaluating the transition in flight-related morphology underlying the origin of early pterygotes.

Wake, 1993), as seen in the reduction of wing venation in partial wings. The number of primary radial veins was correlated with Q and p_w_ (**Fig. 8B**). In other words, large and small phasmids with similar Q tend to have same number of radial veins. Nonetheless, our results further suggest that partial wings are not simply vestigial organs, but possess features potentially influenced by flight-related selection despite high wing loading. Below, we discuss the transition in wing morphology, within a context of how these features may be relevant to partial-wings interacting with upward flow field during descent.

Despite a generally round shape, phasmid wings show reduced aspect ratio with increasing size, albeit following sex-specific trends (**Fig. 7**). Wings of female phasmids showed more complex, stepwise patterns. Why was this only found in female insects? Female phasmids are heavier than males (Zeng et al. 2020), and thus have a greater demand for vertical force generation; expansion of the wing membrane is thus preferred in high p_w_ regime. Also, a low AR wing may help in drag production in gliding (Ennos, 1989).

A heavier phasmid would require a greater number of radial veins to support increased wing area. The most direct evidence for this is that female phasmids exhibit more radial veins than males with the same wing loading (**Fig. 8C**), likely because females are heavier than males at the same body length (Zeng et al., 2020). Demand for increased bending stiffness was also shown by the conservation of AA veins, which support the anterior portion of membrane and are directly associated with wing depressor muscles (**Fig. 8**; **SI Fig. S2**). Within low to median values of p_w_ (∼10 Nm^-2^), these veins likely play important roles in generating downstroke forces, or at least maintaining a relatively stable wing against upward airflow. Further, our observations suggest that the secondary intercalary veins at the wing margin of partial wings likely contribute to an umbrella-like camber with reinforced stiffness (**SI Fig. S3**), as observed in locust wings (Herbert et al., 2000). At the high-p_w_ regime, unlike many miniaturized wings serving non-flight functions, maintenance of the complete wing membrane (as in *Asceles tanarata tanarata*; **Fig. 9A**) suggests reduced yet non-trivial aerodynamic utility.

In addition to generally linear scaling relationships, several parameters exhibited a step-wise transition across p_w_ ≈ 10 Nm^-2^, including wing aspect ratio in females (**Fig. 7B-C**), sharp reduction in the number of anterior anal veins (**Fig. 8D**) and the reduction of secondary radial veins (**Fig. 9**). The sharp reduction in the number of AA veins is perhaps coupled with a major reduction of wingbase stiffness (e.g., the aerodynamic moment exceeds what can be sustained by muscles and the wingbase). These step-wise changes suggest a potentially continuous transition in wing performance, followed by a major relaxation in selective pressure once a threshold of aerodynamic loading is surpassed (see Zeng et al., 2020).

### 4.3 Implications for flight transition in phasmids

Our results provide a suggestive overview of the interactions between wings and the body-leg system in flight. Wing mass and flight musculature are coupled with a transition in wing loading (**Fig. 7**), suggesting that larger wings are powered by stronger flight musculature. Such coupling between wing size and flight musculature is seen in other insects expressing partial wings (e.g., dispersal polymorphism in orthopterans; Zera and Denno, 1997). We also speculate that the reduction in flight muscle also involves reconfiguration of different muscle groups, because partial wing may flap differently (e.g., more power production during the downstroke).

The flight-related morphology of the body-leg system (e.g., center of mass and projected planform area) may have little influenced by flight-related selection (**Fig. 11-12**), suggesting that variation in forces and moments produced by different sized wings would be mechanistically linked to a relatively conserved body-leg inertia. As the abdominal moment of inertia is fairly constant and invariant with respect to p_w_ (**SI Datasheet S2**), inertial moments would be largely dependent on wing movement., Nevertheless, the body-leg system likely plays different aerodynamic roles throughout flight transition. For example, more drag production on the body-leg system (and correspondingly faster descent) would be necessary to offset most body weight in either gliding or parachuting of partial winged species.

At the whole-insect level, further work is needed to address the relationship between forces and moments generated by wings and the body-leg system. With reduced wing forces, partial winged phasmids typically descend during flight, which may benefit from the posterior shift of pterothoracic segments (**Fig. 12F**). This shift leads to a reduced moment arm deriving from wing forces, and thus may benefit passive stability by reducing the wing moment, especially in phasmids with high wing loading and potentially falling with greater speed. This may also have underpinned the elongation of the mesothorax that associates with twig mimicry (Yang et al., 2022).

With little known about the aerodynamics of partial and low aspect ratio wings operating at high p_w_, kinematic data (e.g., flow speed, wing flapping kinematics, and wing deformation) along with force analysis would be necessary to further address functional significance of these features. To understand why different phasmids adopt different body-leg movements in flight, the aerodynamic and inertial roles of body-leg sections should also be analyzed in relation to the dynamics of wing forces and moments. Lastly, phasmids’ flight transition via descent is highly influenced by their generally arboreal life style; nevertheless, environmentally-mediated wing and flight transitions are not rare, as seen in other orthopteroid insects (Braun, 2011; McCulloch et al., 2018).

### 4.4 Implications for flight transition in early pterygotes

The earlies pterygotes show distinct flight-related morphology unseen in modern insects, such as long slender wings resembling those of dragonflies, long cerci resembling those of mayflies and relatively low wing loading (Wootton and Kukalová-Peck, 2000; **Fig. 5-6**). Lacking fossils of intermediate stages, reconstruction of the early flight transition should be hypothesis-based and with morphological data carefully incorporated. Using the multi-dimensional perspective on phasmid flight transition, we suggest that modeling transitional intermediates can be done in two steps (**Fig. 13B**).

First, establishing the wing loading landscape specific to early pterygotes so that possible evolutionary trajectories can be described with respect to L, Q and p_w_. To achieve this, power-law scaling models involving shape and sizes (e.g., wing size and shape; body shape) can be generated using fossil data, and body mass allometry may be inferred from extant insects.

Second, flight-related morphology throughout the transition may be reconstructed using empirical models as references. Throughout initial wing augmentation, there were likely increase of wing aspect ratio, along with an increase in mass and vein reorganization. Also, there were possible shifts in the positions of pterothoracic segments during the gain of flight, and possible size differentiation between the initially three wing pairs. Ancestral pterygotes likely possessed multiple wing pairs, including three thoracic wing pairs along with abdominal wing pads (Haug et al., 2016; Prokop et al. 2022), and all should be considered to determine the longitudinal position of the center of aerodynamic pressure for the wing-body system.

With reconstructed intermediate morphologies, flight mechanisms can then be evaluated using computational and physical models. What cannot be easily solved by modeling is the flapping motion of various sized wings, and the behavioral coordination between moving wings and the body-appendage system. Therefore, having kinematic data (e.g., body speed and wing kinematics) for modern taxa undergoing flight transition would help reconstruction of the process of flight gain through a series of morphological intermediates.

**Acknowledgments:** We thank Francis Seow-Choen for helping with field work and commenting on stick insect natural history. We also thank Steve Yanoviak for sharing data, thank David Wake, Kipling Will, Rosie Gillespie, George Roderick, Paul Brock, Richard Wassersug, two anonymous reviewers and Animal Flight Lab at UC-Berkeley for comments, and thank Datuk Monica Chia, Faye Pon, Joan Chen, Xiaolin Chen, Ho-Yeon Han, Ian Abercrombie, Yamai Shi-Fu Huang, Lin Cao, Azuan Aziz, and Juhaida Harun for help with data collection. We further thank the Sabah Parks and the Forestry Department of Pahang, Malaysia, for permission to collect insects. This research was supported by National Science Foundation (DDIG-1110855), the Museum of Vertebrate Zoology and the Department of Integrative Biology at UC-Berkeley, the Undergraduate Research Apprentice Program (URAP) of UC-Berkeley, the Society for Integrative and Comparative Biology (SICB), and the Ministry of Higher Education Malaysia [FRGS/1/2012/SG03/UKM/03/1(STWN)].

**Author Contributions:** Y.Z. devised experiments and performed the majority of data collection and analyses. S.P., C.G. and S.Y. contributed to data collection and morphological analyses, and F.H. contributed to field data collection. Y.Z. and R.D. wrote the manuscript.

**Conflicts of Interests**: The authors declare no conflicts of interests.

**Data Availability**: The authors confirm that the data supporting the findings of this study are available within the article and its supplementary materials.

## Supplementary Datasheets

S1. Morphological datasets 1-5 for comparisons between different phasmids S2. Results of correlational analyses (PGLS and GLS)

S3. Morphological data of the *Asceles tanarata* species group S4. Morphological data of other pterygotes for comparison

## Symbols and abbreviations

𝑚: insect mass
𝐿: body length
𝐿_𝑤_: wing length
𝐴_𝑤_: single hind wing area
𝑄: = 𝐿_𝑤_/𝐿, relative wing size
𝑝_𝑤_: wing loading
AR: aspect ration
𝐶_𝑤_: wing chord size
𝑁_𝑉_: number of radial veins
AA: anterior anal veins
PA: posterior anal veins
𝜎: = 𝑁_𝑣_/𝐴_𝑤_, mean area density of radial veins
𝑚_𝑤_: total mass (of two hindwings combined)
𝑚^′^: = 𝑚_𝑤_/^𝑚^, relative wing mass
𝑚^′^_fm_: relative mass of flight muscle
*l̂_CM_*: longitudinal position of body-leg center of mass
*l̂_mj_*: longitudinal position of the median joint
*l̂_wb_*: longitudinal position of the wingbase
*r̂_wb_*: the radius from center of mass to wingbase
*r̂_mj_*: the radius from center of mass to median joint

## References

1. Abràmoff, M. D., Magalhães, P. J. and Ram, S. (2004). Image processing with ImageJ. Biophotonics international 11, 36–43.

2. Alexander, D. E. (2018). A century and a half of research on the evolution of insect flight. Arthropod Struct Dev 47, 322–327.

3. Bank, S., Buckley, T. R., Büscher, T. H., Bresseel, J., Constant, J., De Haan, M., Dittmar, D., Dräger, H., Kahar, R. S. and Kang, A. (2021). Reconstructing the nonadaptive radiation of an ancient lineage of ground-dwelling stick insects (Phasmatodea: Heteropterygidae). Systematic Entomology 46, 487–507.

4. Bank, S. and Bradler, S. (2022). A second view on the evolution of flight in stick and leaf insects (Phasmatodea). BMC ecology and evolution 22, 1–17.

5. Bhat, S. S., Zhao, J., Sheridan, J., Hourigan, K. and Thompson, M. C. (2019). Aspect ratio studies on insect wings. Physics of Fluids 31, 121301.

6. Boisseau, R. P., Büscher, T. H., Klawitter, L. J., Gorb, S. N., Emlen, D. J. and Tobalske, B. W. (2022). Multi-modal locomotor costs favor smaller males in a sexually dimorphic leaf-mimicking insect. BMC ecology and evolution 22, 1–18.

7. Bragg, P. E. (1997). A glossary of terms used to describe phasmids. Phasmid Studies 6, 24–33.

8. Brauckmann, C. and Gröning, E. (2018). A reconstruction of Lithomantis varius from Hagen-Vorhalle (Insecta: Palaeodictyoptera: Lithomantidae; early Pennsylvanian, Late Carboniferous, Germany). Entomologia Generalis 37, 231–241.

9. Braun, H. (2011). A brief revision of brachypterous Phaneropterinae of the tropical Andes (Orthoptera, Tettigoniidae, Odonturini). Zootaxa 2991, 35–43.

10. Brock, P. D. (1999). Stick and leaf insects of Peninsular Malaysia and Singapore, Malaysian Nature Society.

11. Carpenter, F. M. (1951). Studies on Carboniferous insects from Commentry, France: Part II. The Megasecoptera. Journal of Paleontology , 336–355.

12. Chang, S. -K., Lai, Y. -H., Lin, Y. -J. and Yang, J. -T. (2020). Enhanced lift and thrust via the translational motion between the thorax-abdomen node and the center of mass of a butterfly with a constructive abdominal oscillation. Physical Review E 102, 062407.

13. Combes, S. A. and Daniel, T. L. (2003). Flexural stiffness in insect wings I. Scaling and the influence of wing venation. Journal of experimental biology 206, 2979–2987.

14. Dirks, J. -H. and Taylor, D. (2012). Veins improve fracture toughness of insect wings.

15. Dudley, R. (2002). The biomechanics of insect flight: form, function, evolution, Princeton Univ Pr.

16. Dudley, R., Byrnes, G., Yanoviak, S. P., Borrell, B., Brown, R. M. and McGuire, J. A. (2007). Gliding and the Functional Origins of Flight: Biomechanical Novelty or Necessity? Annual Review of Ecology, Evolution, and Systematics 38, 179–201.

17. Dudley, R. and Yanoviak, S. P. (2011). Animal aloft: the origins of aerial behavior and flight. Integr Comp Biol 51, 926–936.

18. Dyhr, J. P., Morgansen, K. A., Daniel, T. L. and Cowan, N. J. (2013). Flexible strategies for flight control: an active role for the abdomen. Journal of Experimental Biology 216, 1523–1536.

19. Ellington, C. P. (1984). The aerodynamics of hovering insect flight. II. Morphological parameters. Philosophical Transactions of the Royal Society of London. Series B, Biological Sciences (1934-1990) 305, 17-40.

20. Ellington, C. P. (1991). Aerodynamics and the origin of insect flight. Adv. Insect Physiol 23, 171–210.

21. Ennos, A. R. (1989). The effect of size on the optimal shapes of gliding insects and seeds. Journal of Zoology 219, 61–69.

22. Forni, G., Martelossi, J., Valero, P., Hennemann, F. H., Conle, O., Luchetti, A. and Mantovani, B. (2022). Macroevolutionary Analyses Provide New Evidences of Phasmid Wings Evolution as a Reversible Process. Syst Biol .

23. Hanken, J. and Wake, D. B. (1993). Miniaturization of body size: organismal consequences and evolutionary significance. Annual Review of Ecology and Systematics 24, 501–519.

24. Haug, J. T., Haug, C. and Garwood, R. J. (2016). Evolution of insect wings and development--new details from P alaeozoic nymphs. Biological Reviews 91, 53–69.

25. Heidinger, I. M. M., Hein, S., Feldhaar, H. and Poethke, H. -J. (2018). Biased dispersal of *Metrioptera bicolor*, a wing dimorphic bush-cricket. Insect science 25, 297–308.

26. Herbert, R. C., Young, P. G., Smith, C. W., Wootton, R. J. and Evans, K. E. (2000). The hind wing of the desert locust (Schistocerca gregaria Forskal). III. A finite element analysis of a deployable structure. Journal of Experimental Biology 203, 2945–2955.

27. Kruyt, J. W., Van Heijst, G. F., Altshuler, D. L. and Lentink, D. (2015). Power reduction and the radial limit of stall delay in revolving wings of different aspect ratio. Journal of the Royal Society Interface 12, 20150051.

28. Kukalova-Peck, J. (1971). The structure of Dunbaria (Palaeodictyoptera). Psyche 78, 306–318.

29. Kukalová-Peck, J. (1983). Origin of the insect wing and wing articulation from the arthropodan leg. Can J Zool 61, 1618–1669.

30. Le Roy, C., Debat, V. and Llaurens, V. (2019). Adaptive evolution of butterfly wing shape: from morphology to behaviour. Biological Reviews 94, 1261–1281.

31. Lehmann, F. -O. and Dickinson, M. H. (1997). The changes in power requirements and muscle efficiency during elevated force production in the fruit fly *Drosophila melanogaster*. J Exp Biol 200, 1133–1143.

32. Mani, M. S. (2013). Ecology and biogeography of high altitude insects, Springer Science & Business Media.

33. Marden, J. H. (1987). Maximum lift production during takeoff in flying animals. Journal of experimental Biology 130, 235–258.

34. Marden, J. H. (2003). The surface-skimming hypothesis for the evolution of insect flight. Acta zoologica cracoviensia 46, 73–84.

35. McCulloch, G. A. and Waters, J. M. (2018). Does wing reduction influence the relationship between altitude and insect body size? A case study using New Zealand’s diverse stonefly fauna. Ecol Evol 8, 953–960.

36. Orme, D., Freckleton, R., Thomas, G. and Petzoldt, T. (2013). The caper package: comparative analysis of phylogenetics and evolution in R. R package version 5, 1–36.

37. Paradis, E., Claude, J. and Strimmer, K. (2004). APE: analyses of phylogenetics and evolution in R language. Bioinformatics 20, 289–290.

38. Pecharova, M., Ren, D. and Prokop, J. (2015). A new palaeodictyopteroid (Megasecoptera: Brodiopteridae) from the Early Pennsylvanian of northern China reveals unique morphological traits and intra-specific variability. Alcheringa: An Australasian Journal of Palaeontology 39, 236–249.

39. Prokop, J. and Engel, M. S. (2019). Palaeodictyopterida. Current Biology 29, 306–309.

40. Prokop, J., Krzemińska, E., Krzemiński, W., Rosová, K., Pecharová, M., Nel, A. and Engel, M. S. (2019). Ecomorphological diversification of the Late Palaeozoic Palaeodictyopterida reveals different larval strategies and amphibious lifestyle in adults. R Soc Open Sci 6, 190460.

41. Prokop, J., Rosová, K., Krzemińska, E., Krzemiński, W., Nel, A. and Engel, M. S. (2022). Abdominal serial homologues of wings in Paleozoic insects. Current Biology 32, 3414–3422.

42. Ragge, D. R. (1955). The wing-venation of the order Phasmida. Transactions of the Royal entomological Society of London 106, 375–392.

43. Ritz, C., Baty, F., Streibig, J. C. and Gerhard, D. (2015). Dose-Response Analysis Using R. PLoS One 10.

44. Roff, D. A. (1994). The evolution of flightlessness: is history important? Evolutionary Ecology 8, 639–657.

45. Schachat, S. R., Goldstein, P. Z., Desalle, R., Bobo, D. M., Boyce, C. K., Payne, J. L. and Labandeira, C. C. (2023). Illusion of flight? Absence, evidence and the age of winged insects. Biological Journal of the Linnean Society 138, 143–168.

46. Seow-Choen, F. (2000). An illustrated guide to the stick and leaf insects of Peninsular Malaysia and Singapore, Natural History Publications (Borneo).

47. Smart, J. (1953). The wing-venation of the migratory locust (*Locusta migratoria* Linn.) (Insecta: Acridiidae). Proceedings of the Zoological Society of London 123, 207–217.

48. Stevenson, R. D., Hill, M. F. and Bryant, P. J. (1995). Organ and cell allometry in Hawaiian *Drosophila*: how to make a big fly. Proceedings of the Royal Society of London. Series B: Biological Sciences 259, 105–110.

49. Sunada, S., Yasuda, T., Yasuda, K. and Kawachi, K. (2002). Comparison of wing characteristics at an ultralow Reynolds number. Journal of aircraft 39, 331–338.

50. Thomas, A. L. and Taylor, G. K. (2001). Animal flight dynamics I. Stability in gliding flight. J Theor Biol 212, 399–424.

51. Taylor, G. K. and Thomas, A. L. (2003). Dynamic flight stability in the desert locust Schistocerca gregaria. Journal of Experimental Biology 206, 2803–2829.

52. Torres, G. E. and Mueller, T. J. (2004). Low-aspect-ratio wing aerodynamics at low Reynolds number. AIAA Journal 42, 865–873.

53. Wootton, K. J. and Kukalová-Peck, J. (2000). Flight adaptations in Palaeozoic Palaeoptera (Insecta). Biol Rev Camb Philos Soc 75, 129–167.

54. Yang, H., Engel, M. S., Zhang, W., Ren, D. and Gao, T. (2022). Mesozoic insect fossils reveal the early evolution of twig mimicry. Science bulletin 67, 1641–1643.

55. Zeng, Y., Lam, K., Chen, Y., Gong, M., Xu, Z. and Dudley, R. (2017). Biomechanics of aerial righting in wingless nymphal stick insects. Interface Focus 7, 20160075.

56. Zeng, Y., O’Malley, C., Singhal, S., Rahim, F., Park, S., Chen, X. and Dudley, R. (2020). A tale of winglets: evolution of flight morphology in stick insects. Frontiers in Ecology and Evolution 8, 121.

57. Zera, A. J. and Denno, R. F. (1997). Physiology and ecology of dispersal polymorphism in insects. Annual review of entomology 42, 207–230.

