## Supplementary Information for "Beyond winglets: evolutionary scaling of flight-related morphology in stick insects (Phasmatodea)"

### Supplementary Text

#### A. Environmentally associated body and wing reduction in *A. tanarata*

Evolutionary coupling between wing and body reduction within *A. tanarata* is distinct from the more general pattern in stick insects, for which wing and body sizes tend to be inversely correlated (e.g., larger females possess smaller wings; **SI Fig. S4A,B**). Below, we examine the patterns of morphological reduction in *A. tanarata* and their underlying processes.

For simplicity, the six study morphs within *A. tanarata* are coded numerically in order of increasing altitudinal occurrence (i.e., *A. tanarata singapura*, At1; *A. tanarata amplior*, At2; *A. tanarata tanarata*, At3), and with the suffixes 'f' and 'm' denoting 'female' and 'male', respectively.

##### A.1 Body size reduction and mass distribution

From the lowlands to the highlands, body length declined by ~46% in females and by ~37% in males. Similarly, body mass reduction was ~72% in females and ~65% in males. Nevertheless, these reductions followed very similar power-law scaling with body length, with the mean scaling exponent $a$ ~2.2 in both sexes (**SI Fig. 4C**). Mass proportions of the anterior and posterior sections showed no significant correlation with body mass across three subspecies (head and thorax, 23% – 25% reduction in males and 18% – 22% females; abdomen, 41% – 46% reduction in males and ~66% in females) (**SI Fig. S4D**). However, there was a significant increase in the mass percentage of the legs in males, increasing from 17% to 24%, and a corresponding increased relative length of mid-legs in both sexes (**SI Fig. 45E**). Trending in the opposite direction, the mass percentage of thoracic musculature decreased from 14% to 5% in males and from 7% to 4% in females; wing mass declined from 2.2% – 2.8% to <1% (**SI Fig. S4D**).

Mass allometry was similar for both sexes, although the specific mechanism associated with environmental change at higher elevation is unclear. From sea level to 1600 m, oxygen availability and atmospheric pressure reduce by ~20% (Mani, 2013), and there is also substantial variation in air temperature, seasonality, and precipitation among the home localities of each subspecies (see **SI Fig. S7**).

Growth patterns in *A. tanarata* may be influenced by such changes, although knowledge of life history (e.g., growth period and relationship with host plants) are not well known. One possible cause is a shortened growth cycle at lower air temperatures, as reported for some Orthoptera (Sømme, 1989). Body size reduction may also associate with altitudinal variation in forest composition (e.g., Pendry and Proctor, 1996; Proctor et al., 1988), which in turn may influence growth. For example, *Asceles* females pin eggs to the leaves of host plants (Sellick, 1993), suggesting relatively less demand for dispersal in newly hatched nymphs as compared to other phasmid species (e.g., *Extatosoma tiaratum*; see Zeng et al., 2020b).

Selection for increased female fecundity may in part counteract body size reduction, although such an effect was not observed here given the similar mass allometry between the two sexes. However, relative egg mass (i.e., the ratio of egg mass to insect mass) did increase in females as body size decreased, even though individual eggs became smaller (**SI Fig. S4F,G**). Females at higher elevation may thus lay fewer but relatively larger eggs (see also Shapiro, 1986). Overall, these results suggest powerful effects of selection on this stick insect lineage, although the altitudinal range in question is much reduced relative to that commonly referred to as the montane environment (e.g., > 2000 m; Mani, 2013).

##### A.2 Linear trajectories of wing loading variation

The altitudinal increase of wing loading ($p_w$) showed sex-specific and linear trends, with females exhibiting greater $p_w$ at all altitudes. In females, $p_w$ increased by a factor of ~5 (from ~6.4 to ~30.0 $Nm^{-2}$); in males, $p_w$ increased about 10-fold (from ~2.0 to ~24.3 $Nm^{-2}$) (**SI Fig. S5**). In the lowland At1, $p_w$ of females was ~3.2 times greater than that of the male, but this ratio declined to ~1.2 in the highland At3. Overall, both sexes converged to similarly high values of $p_w$. The large increase in wing loading can be represented by the scaling equation from Zeng et al. (2020a): $p_w = \frac{mg}{A_w} \propto$

$L^{a-b}Q^{-b}$, wherein the power-law exponent of body mass $a$ is ~2.2 (**SI Fig. S4C**), and the power-law exponent of the ratio of wing area to wing length $b$ is either ~1.8 (males) or ~2.3 (females). The sex-specific scaling of $p_w$ can then be represented as $p_w \propto L^{-0.1}Q^{-2.3}$ (females) and as $p_w \propto L^{0.4}Q^{-1.8}$ (males).

##### A.3 Wing reduction

With increasing altitude, males showed a transition from long wings (relative wing size Q ≈ 0.59) to miniaturized wings (Q ≈ 0.16), whereas females showed a transition from intermediate-sized (Q ≈ 0.44) to miniaturized wings (Q ≈ 0.25) (**SI Fig. S4B**). Wing reduction followed sex-specific trends with respect to body size, whereby the reduction in relative wing size Q was steeper in males. Comparing At1 to At3, the total reduction in Q was ~70% in males and ~40% in females, corresponding to ~97% and ~94% absolute reduction in wing area, respectively. The reduction of AR over the $p_w$ range of 4 – 14 $Nm^{-2}$ (At1f and both sexes of At2) is coupled with an increase in the relative area of wing membrane. In both sexes, the areal percentage of wing membrane ($A_{mem}$) dropped from ~85% to ~70%. Over the $p_w$ range of 4 – 14 $Nm^{-2}$, $A_{mem}$ is 3% – 5% greater than what expected from a linear reduction (**SI Fig. S6B**).

##### A.4 Flight evolution in *A. tanarata*

Wing reduction follows sex-specific trajectories but more generally may reflect developmental truncation as seen in other high-altitude insects (e.g., stoneflies; McCulloch et al., 2016), which is different from a flight-reproduction tradeoff (e.g., as in wing-dimorphic crickets; Mole and Zera, 1993; Guerra, 2011). Such developmental truncation may also underlie high-altitude wing reduction in other sympatric phasmid species (e.g., *Lopaphus* spp. and *A. marginatus*; see Seow-Choen, 2000; Bragg, 2001).

Unlike other montane insects with reduced wings and living in treeless habitats (see Roff, 1994; Dillon et al., 2006), the *A. tanarata* subspecies studied here lives in well-developed vegetational canopies at all localities (e.g., canopy heights of ~15 m, ~10 m, and ~5 m for At1, At2 and At3, respectively; Seow-Choen, 2000). Selection on aerial ability may be important here, as even wingless larvae of stick insects exhibit directed aerial descent when falling from heights (see Zeng et al., 2020b). By contrast, some sympatric phasmid species found along the same altitudinal gradient exhibit neither wing reduction nor flight loss (e.g., *Necroscia punctata* and *Heterpteryx dilatata*; Seow-Choen, 2000; YZ, pers. observ.). Additionally, the substantial wing membrane of the miniaturized wings of At3 (**Fig. 9B**) suggests aerodynamic utility during descent (e.g., in aerial escape; Bedford, 1978). Detailed behavioral and phylogenetic studies of phasmid wing use during falls and glides are necessary to evaluate their actual biomechanical function and how they may have evolved.

##### A.5 Summary

Our analyses suggest selection for body size-reduction acting on both sexes, suggesting a dwarfism-induced developmental truncation of wings and flight-related structures. Although body size reduction theoretically contributes to a reduction in wing loading, wing size reduction in *A. tanarata* is predominantly associated with an increase in wing loading.

#### Supplementary Figures

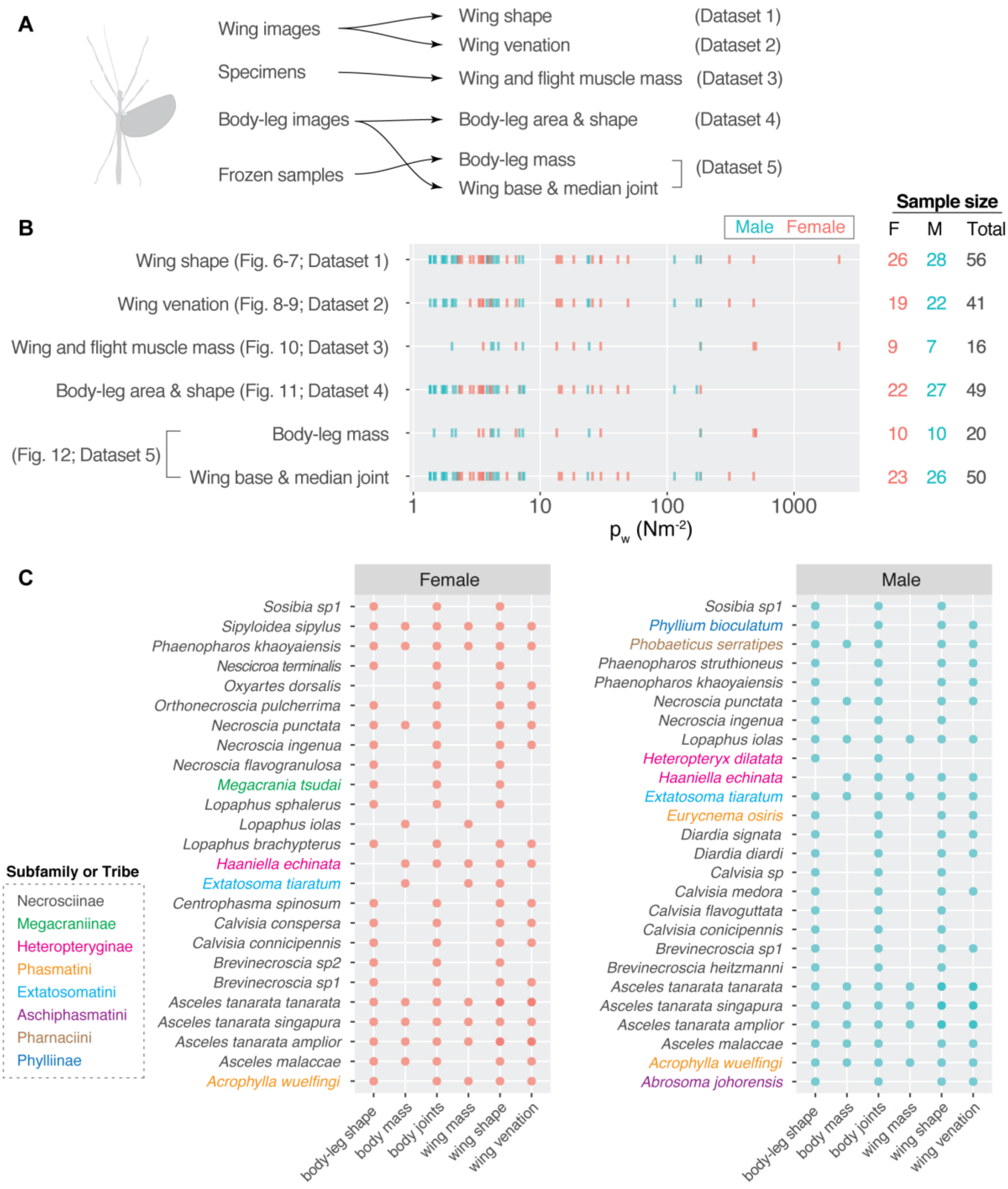


**Figure S1. Summary of workflow and phasmid samples.**
**(A)** Data processing workflow for each of the five main datasets. For Dataset 4 (body-leg area and shape), the leaf insect *Phyllium bioculatum* was excluded. **(B)** A summary of coverage of wing loading for each dataset; sample sizes are annotated to the right. **(C)** A summary of representation for each sample (species + sex) in each dataset. Colors denote different subfamilies or tribes, whereby the majority of samples were members of the subfamily Necrosciinae.

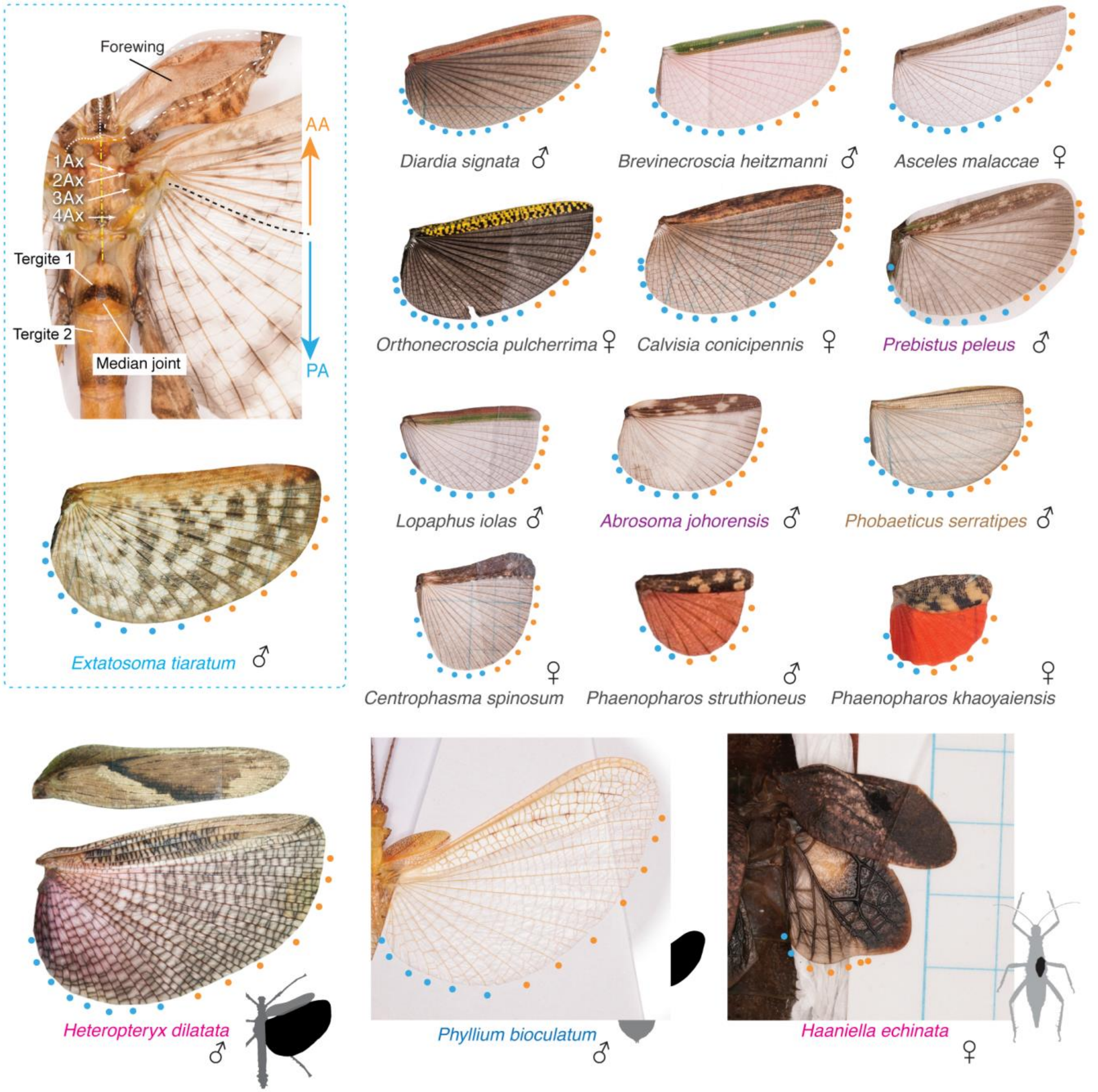


**Fig. S2. Gallery of sampled phasmid wings.**
Colors of species names represent either subfamilies or tribes. Thoracic morphology is demonstrated with an *Extatosoma tiaratum* male. The AA and PA vein groups can be clearly distinguished from the close association between AA veins and the 3rd axillary sclerite (3Ax). Median joint, the joint between 1st and 2nd abdominal tergum. We highlight two cases of less condensed costal edges: (1),wings of male *Phyllium bioculatum*, which cover a flat, laterally expanded abdomen, and (2), wings of female *Haaniella echinata*, which cover a wide abdomen and also function as a stridulatory organ. Note *Prebistus peleus* (Aschiphasmatini) was not included in main datasets due to the lack of wing loading data.

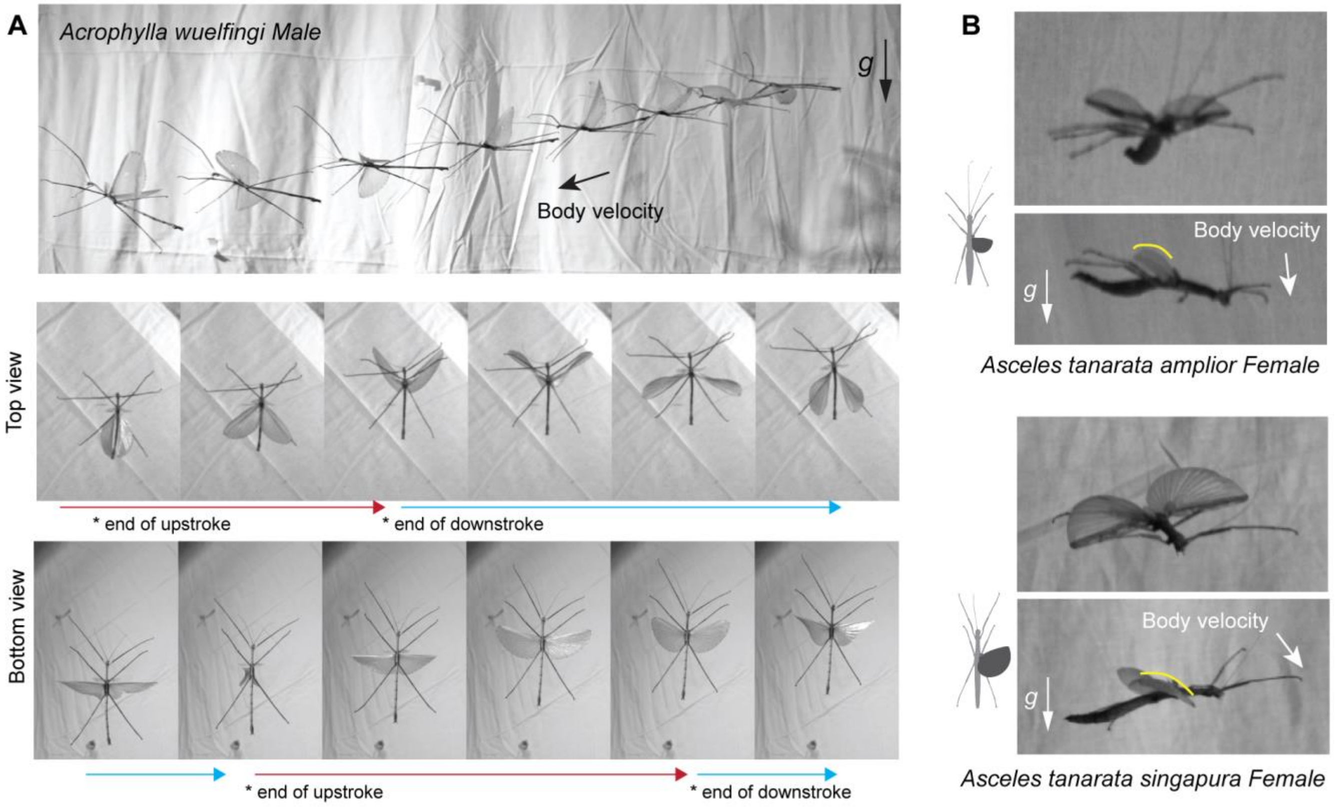


**Figure S3. Observation of wing deformation in fully-winged and partial-winged phasmids, as captured with high-speed video (250 fps).**

**(A)** Sample sequences from flight trials for a long-winged phasmid (*Acrophylla wuelfingi* male; L ~13 cm, $p_w$ ~4.3 $Nm^{-2}$). Note the dynamic wing deformations, including a 'clap and peel' wing motion during the initiation of the downstroke. **(B)** Wing camber (indicated with yellow lines) in two partial-winged phasmids (*Asceles tanarata amplior* female: L ~5 cm, $p_w$ ~13.5 $Nm^{-2}$; *Asceles tanarata singapura* female; L ~7 cm, $p_w$ ~6.4 $Nm^{-2}$).

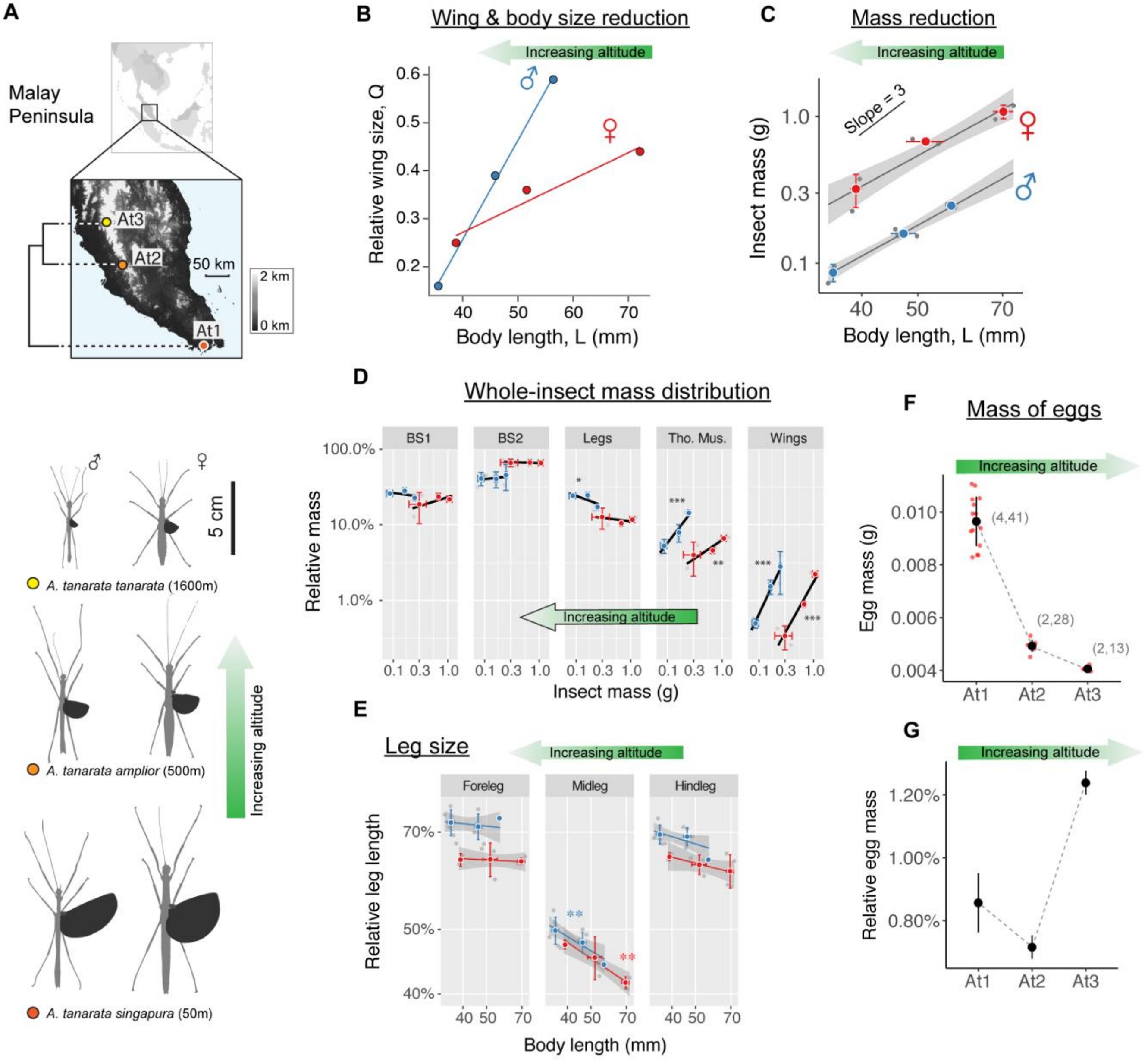


**Figure S4. Altitudinal variation of morphology in *Asceles tanarata* species group.**

**(A)** Dorsally projected profiles of the three subspecies of *A. tanarata*, showing reductions in body and wing size at greater altitude. On the left is an altitudinal map of the Malay Peninsula, annotated with locations where three subspecies were collected; at right are the phylogenetic relationships among three subspecies, with nodes annotated by divergence time (see Zeng et al., 2020a). **(B)** Sex-specific trends in wing and body reduction, as shown by variation in relative wing size (Q) versus body size (L). **(C)** Power-law scaling of insect mass with respect to body length. Mass allometries for male and female insects showed similar coefficients (females, 2.2±0.3; males, 2.2±0.2; mean±s.e.m.). Trend lines represent linear regression models, with shading representing s.e.m. **(D)** Comparison of masses of different body parts with respect to whole-insect mass. Values correspond to means±S.D., with black trend lines based on generalized least-square linear regression. Asterisks represent significance of correlation (*, $P < 0.05$; **, $P < 0.01$; ***, $P < 0.001$). BS1, head and thorax; BS2, abdomen; Tho. Mus., pterothoracic musculature. Sample sizes: At1m and At2f, N = 2; At1f, At2m and At3f, N = 3; At3m, N = 5.

**(E)** Relative length of midlegs increases with increasing altitude. The plot compares leg lengths with respect to body length on log-transformed axes. Values correspond to mean±S.D., with trend lines based on generalized least-square linear regression. Asterisks represent significance of correlation (**, $P < 0.01$). Red, females; blue, males. Sample sizes: At1m and At2f, N = 2; At1f, At2m and At3f, N = 3; At3m, N = 5. **(F)** Egg mass for the three subspecies, showing reduction of ~60% from lowland At1 to highland At3. Values represent mean±S.D., with the number of sampled individuals and total number of eggs in brackets. Egg mass was estimated from groups of 3 – 4 eggs using a portable electronic balance (PP-2060D, Acculab; accuracy 0.001 g); the mean from each group is represented by red dots. **(G)** Relative egg mass is reduced in At2 but is substantially greater in At3 from Genting Highland. Based on two-way ANOVA post-hoc pairwise comparisons, the relative egg mass was significantly different between all three subspecies ($P < 0.0001$).

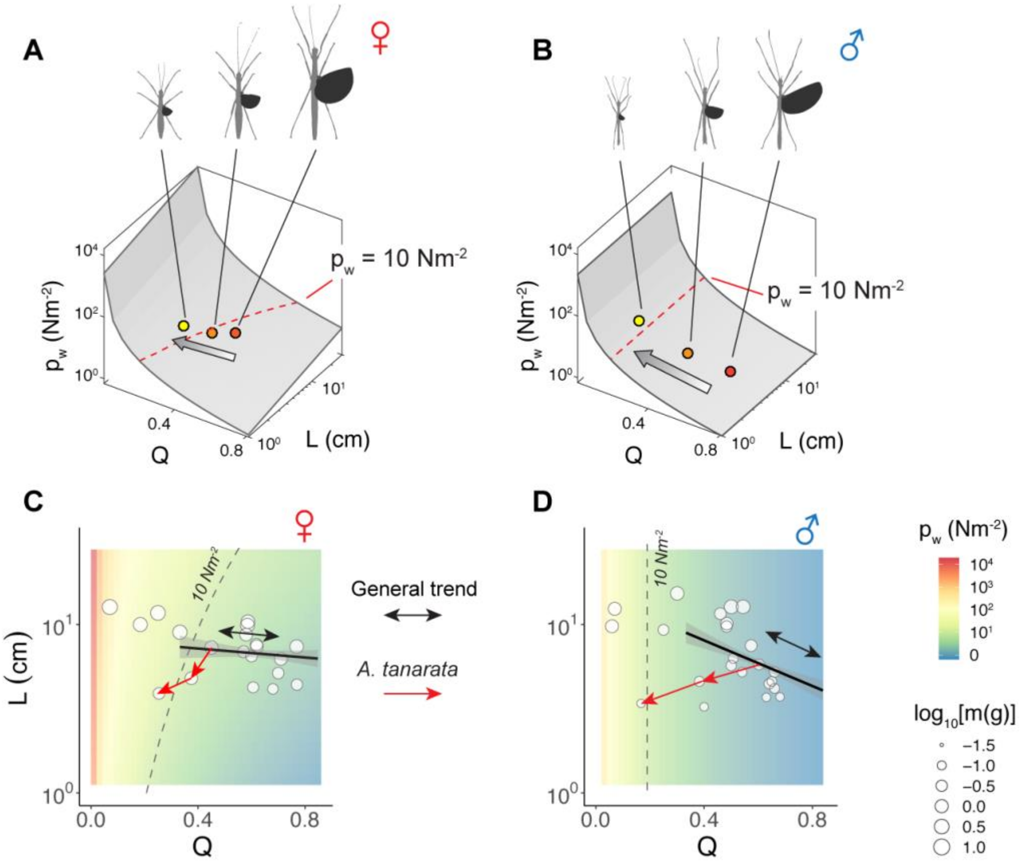


**Figure S5. Evolution of wing loading in *Asceles tanarata* species group.**

**(A)-(B)** Evolutionary trajectories of *A. tanarata* on the wing loading ($p_w$) landscapes of male and female phasmids. Note sex-specific topologies of the $p_w$ landscape and trajectories of morphological evolution. Arrows represent the direction of increasing altitude. Red dashed lines representing a threshold of $p_w$ = 10 Nm$^{-2}$ for flight loss (see Zeng et al., 2020a). **(C)** Female and **(D)** male phasmids sampled in this study and plotted on sex-specific $p_w$ landscapes, with dot size representing insect mass. Solid trend lines represent the general pattern in flight transition – an inverse correlation between L and Q in flight evolution (based on ~270 morphologies for each sex; see Zeng et al., 2020a). As an outlier, the *A. tanarata* species group (annotated with red arrows) exhibited intraspecific reductions in both L and Q with increasing altitude.

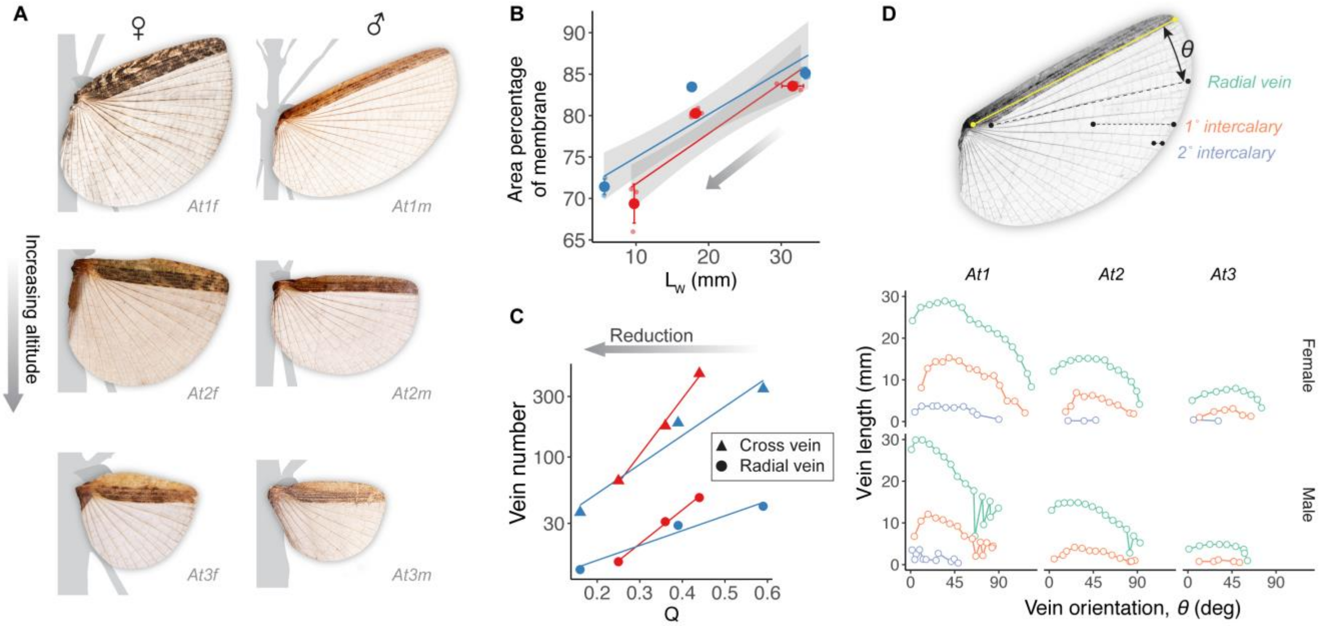


**Figure S6. Variation of wing morphology in *A. tanarata*.**

**(A)** Photographs of fully unfolded wings showing reduction in venation as wing size reduces. The miniaturized wings in At3 retain cross vein components in the membranous region (see also **Fig. 9A,B**). **(B)** Wing size reduction derives from a disproportionately greater loss of the membranous region, as shown by the general linear trend in reduction of the membranous region. There was a clear disparity between an expected linear reduction and the actual value over the intermediate range of wing loading (approximately 4 – 14 $Nm^{-2}$). **(C)** As relative wing size reduces, vein count declines. In females, the ratio of cross-vein to radial-vein number declined from ~9.5 (At1f) to ~4.3 (At3f); in males, this ratio declined from ~8.4 (At1m) to ~2.8 (At3m). Values represent means±S.D., with trend lines based on generalized linear regression; red, females; blue, males. **(D)** The vein orientation angle with respect to the costal edge (θ, demonstrated on the top) was measured on one wing from each of the six flight morphs. Plot shows variation of vein length with respect to vein orientation; note the relatively even angular distance between adjacent veins. The angular distance ranges from 6.8˚ – 8.9˚, 7.6˚ – 10.5˚ and 9.5˚ – 25.3˚ in radial veins, primary intercalary veins, and secondary intercalary veins, respectively.

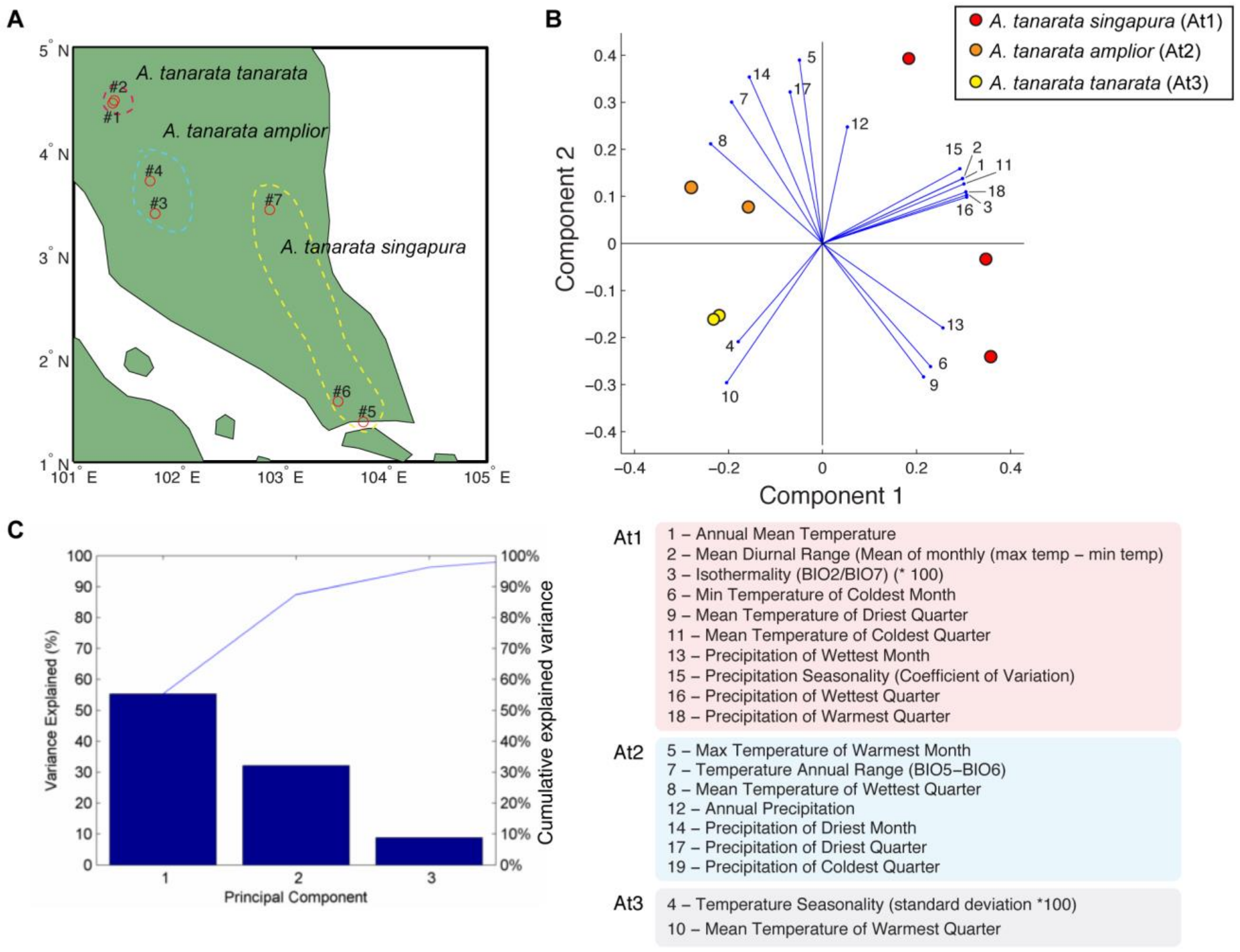


**Figure S7. Climatic variation among locales for the three subspecies.**

**(A)** We first extracted bioclimatic variables for the type localities (Brock, 1999; Seow-Choen, 2000) from the WorldClim database (Hijmans et al., 2005), and then performed a principal component analysis (PCA) in R. **(B)** Subspecies locales showed a distinct clustered distribution, with that of At1 being the most different from those of the other two, mostly in terms of temperature and precipitation cycles (red-highlighted variables in legend). In addition, the highland At3 locality experiences greater seasonality and lower air temperatures (variables 4 and 10) as compared to that of At2. **(C)** Percentage of explained PCA variance, with components 1 and 2 together explaining ~85% of total variance.

#### Supplementary References

**Bedford, G. O.** (1978). Biology and ecology of the Phasmatodea. *Annual Review of Entomology* **23**, 125-149.

**Bragg, P. E.** (2001). *Phasmids of Borneo*, Kota Kinabalu, Sabah: Natural History Pub., 2001.

**Dillon, M. E., Frazier, M. R. and Dudley, R.** (2006). Into thin air: physiology and evolution of alpine insects. *Integrative and Comparative Biology* **46**, 49-61.

**Guerra, P. A.** (2011). Evaluating the life-history trade-off between dispersal capability and reproduction in wing dimorphic insects: a meta-analysis. *Biological Reviews* **86**, 813-835.

**Hijmans, R. J., Cameron, S. E., Parra, J. L., Jones, P. G. and Jarvis, A.** (2005). Very high resolution interpolated climate surfaces for global land areas. *International Journal of Climatology: A Journal of the Royal Meteorological Society* **25**, 1965-1978.

**Mani, M. S.** (2013). *Ecology and biogeography of high altitude insects*, Springer Science & Business Media.

**McCulloch, G. A., Wallis, G. P. and Waters, J. M.** (2016). A time-calibrated phylogeny of southern hemisphere stoneflies: Testing for Gondwanan origins. *Molecular Phylogenetic Evolution* **96**, 150-160.

**Mole, S. and Zera, A. J.** (1993). Differential allocation of resources underlies the dispersal-reproduction trade-off in the wing-dimorphic cricket, *Gryllus rubens*. *Oecologia* **93**, 121-127.

**Pendry, C. A. and Proctor, J.** (1996). The causes of altitudinal zonation of rain forests on Bukit Belalong, Brunei. *Journal of Ecology*, **84**, 407-418.

**Proctor, J., Lee, Y. F., Langley, A. M., Munro, W. R. C. and Nelson, T.** (1988). Ecological studies on Gunung Silam, a small ultrabasic mountain in Sabah, Malaysia. I. Environment, forest structure and floristics. *The Journal of Ecology*, **76**, 320-340.

**Roff, D. A.** (1994). The evolution of flightlessness: is history important? *Evolutionary Ecology* **8**, 639-657.

**Sellick, J.** (1993). The leaf-piercing eggs of *Asceles*. *Phasmid Studies* **2**, 54-55.

**Seow-Choen, F.** (2000). *An illustrated guide to the stick and leaf insects of Peninsular Malaysia and Singapore*, Natural History Publications (Borneo).

**Shapiro, A. M.** (1986). r-K selection at various taxonomic levels in the Pierine butterflies of North and South America. In *The evolution of insect life cycles*, pp. 135-152. Springer.

**Sømme, L.** (1989). Adaptations of terrestrial arthropods to the alpine environment. *Biological Reviews* **64**, 367-407.

**Zeng, Y., O'Malley, C., Singhal, S., Rahim, F., Park, S., Chen, X. and Dudley, R.** (2020a). A tale of winglets: evolution of flight morphology in stick insects. *Frontiers in Ecology and Evolution* **8**, 121.

**Zeng, Y., Chang, S. W., Williams, J. Y., Nguyen, L. Y. -N., Tang, J., Naing, G., Kazi, C. and Dudley, R.** (2020b). Canopy parkour: movement ecology of post-hatch dispersal in a gliding nymphal stick insect, *Extatosoma tiaratum*. *Journal of Experimental Biology* **223**, jeb226266.
